# Virion Proteomics of Genetically Intact HCMV Reveals a Novel Regulator of Envelope Glycoprotein Composition that Protects Against Humoral Immunity

**DOI:** 10.1101/2024.11.29.626001

**Authors:** Kirsten Bentley, Evelina Statkute, Isa Murrell, Ceri A. Fielding, Robin Antrobus, Hannah Preston, Lauren Kerr-Jones, Daniel Cochrane, Ilija Brizic, Paul J. Lehner, Gavin W.G. Wilkinson, Eddie C.Y. Wang, Stephen C. Graham, Michael P. Weekes, Richard J. Stanton

**Affiliations:** Infection and Immunity, Cardiff University School of Medicine, Cardiff, CF14 4XN; Cambridge University Institute for Medical Research, Cambridge University, Cambridge CB2 0XY; Center for Proteomics, School of Medicine, University of Rijeka, Rijeka, Croatia; Cambridge Institute for Therapeutic Immunology and Infectious Disease, Cambridge University, Cambridge; Department of Pathology, University of Cambridge, Cambridge, CB2 1QP

## Abstract

Human cytomegalovirus (HCMV) is a clinically important herpesvirus that has co-evolved for millions of years with its human host, and establishes lifelong persistent infection. A substantial proportion of its 235kb genome is dedicated to manipulating host immunity through targeting antiviral host proteins for degradation or relocalisation. Quantitative proteomics of the infected cell has extensively characterised these processes, but the cell-free virion has been less well studied. We therefore carried out proteomic analysis of a clinical HCMV strain (Merlin) virion. This revealed 18 novel components, including the viral protein gpUL141, which is recognised as an NK immune-evasin that targets several host proteins (CD155, CD112, and TRAILR) when expressed within the cell. Co-Immunoprecipitation of gpUL141 from virions identified interactions with viral entry glycoproteins from the trimer (gH/gL/gO), pentamer (gH/gL/UL128/UL130/UL131A), and gH/gpUL116 complexes, as well as gB. Only interactions with gH/gB occurred in the absence of other viral proteins. Analysis supported a model in which gpUL141 homodimers independently interacted with separate gB/gH-containing complexes. gpUL141 encodes an ER retention domain that restricts trafficking through the ER/golgi, and limited the transport of glycoprotein complexes bound by gpUL141. As a result, gpUL141 reduced levels of multiple glycoprotein complexes on the infected cell surface as well as in the virion. This reduced syncytium formation, inhibited antibody-dependent cellular cytotoxicity (ADCC), and reduced susceptibility to neutralising antibodies. Thus, gpUL141 represents an immune-evasin that not only targets host proteins to limit NK-cell attack, but also alters the trafficking of multiple viral glycoprotein complexes in order to evade humoral immunity.

**Significance:** The large number of immune-modulators encoded by HCMV has led to it becoming a paradigm for pathogen-mediated immune-modulation. These mechanisms have informed on virus pathogenesis, the evolution and function of host defences, and identified therapeutic targets. Previously discovered immune-evasins functioned by modifying host proteins. gpUL141 represents a novel strategy in which multiple immunological functions are targeted through manipulating eight different viral entry glycoproteins, via interactions with two proteins common to multiple complexes. Entry glycoproteins define virus tropism, mechanisms of infection and spread, and susceptibility to neutralising and non-neutralising antibody activities, yet UL141 is frequently mutated in passaged viruses. This demonstrates the need to work with virus strains encoding the complete repertoire of viral accessory proteins during preclinical therapeutic development.

## Introduction

Human cytomegalovirus (HCMV) is a clinically significant betaherpesvirus that establishes lifelong persistence in the majority of people worldwide. It is a significant cause of morbidity and mortality in the immunocompromised, and the leading infectious cause of congenital malformation. To date, no vaccine is licensed, and antivirals are limited by toxicity and the selection of resistance mutations.

HCMV has the largest genome of any human virus and has co-evolved with its human host over millions of years. This has enabled it to develop an exceptionally broad range of mechanisms to manipulate host immunity to promote persistence. Only 41-45 of the 170 canonical ORFs are essential for replication *in vitro*, with the remainder providing accessory functions that are required to promote persistence *in vivo*^1,2^. These include proteins that prevent cell death, inhibit innate and intrinsic immunity, limit T-cell recognition, and prevent NK-cell attack, through targeting host antiviral proteins for degradation or relocalisation^3–5^. As a result of these activities, HCMV has become a paradigm for pathogen-mediated immune-evasion.

A challenge of studying these viral accessory functions in HCMV is that mutations in non-essential genes are invariably selected when clinical viruses are passaged *in vitro*. Although the loss of a 15kb genetic region encoding >20 genes in the AD169 strain is perhaps the most extreme example, even short-term passage rapidly selects for mutations^6^. As a result, laboratory strains differ both genotypically and phenotypically from the causative agent of disease^7^. To address this problem, we previously generated an infectious bacterial artificial chromosome (BAC) clone of a HCMV strain (Merlin) for which we had the sequence of the original clinical material^7,8^. This allowed us to correct *in vitro* acquired mutations and re-generate virus containing a genome that exactly matched the patient sample. Like clinical isolates, this virus also accrued mutations upon passage *in vitro*, with RL13 and a complex of three genes, UL128, UL130, UL131A (the UL128 locus; UL128L), the most rapidly selected. Engineering of the Merlin BAC enabled us to selectively repress these genes when virus stocks were grown *in vitro*, permitting recovery of high titre virus for experimental use without risk of mutation.

Study of strain Merlin has led to the discovery of multiple phenomena which differ between passaged and clinical viruses, including extensive virus-induced modulation of the cell-surface proteome through metalloproteinase manipulation^9^, novel cell-surface therapeutic targets^10^, immune-evasion mediated by actin manipulation^11^, functions that prevent Natural Killer cell attack^12–14^, and mechanisms that promote a form of cell-associated spread that is highly resistant to neutralising antibodies^8,15^. Many of these discoveries were underpinned by extensive use of quantitative proteomics analysis to understand how HCMV manipulates the infected cell to drive viral survival following entry^16–27^. However, the virion itself has not been subject to the same level of investigation. Previous mass spectrometry analyses have been carried out using the passaged AD169^28,29^ and TB40^30,31^ strains. However, given the existence of mutations in these strains, we carried out an analysis of virions derived from the Merlin BAC with the intention of producing a wildtype HCMV virion proteome. We unexpectedly discovered that glycoprotein UL141 (gpUL141) is a novel virion component. Although previously demonstrated to manipulate host ligands (CD155, CD112, and TRAILR) to limit NK activation^12–14^, we now show that it has also been co-opted by HCMV to regulate the trafficking of the majority of viral entry glycoproteins to the cell surface and the virion particle in order to provide resistance to multiple different aspects of humoral immunity.

## Results

### The wildtype HCMV virion contains multiple proteins absent from passaged strains

To define the complete repertoire of proteins associated with the strain Merlin virion, we used mass spectrometry to analyse virions that had been separated from non-infectious particles and dense bodies on glycerol-tartrate gradients. We individually analysed six Merlin variants that we were working with as part of previous studies: a wildtype genome, a genome mutated for RL13, and a genome mutated for both RL13 and UL128L, to assess alterations deriving from the presence or absence of the pentameric complex and/or gpRL13^8^; two viruses were included that contained point mutations in an intron of UL128 that result in intermediate UL128L expression levels enabling comparisons with passaged strains of HCMV that retain pentamer expression but at lower levels^32^; and a virus lacking UL141^13^. In addition to analysing virion preparations individually, quantitative comparisons were made by producing virions in media containing different isotopes of arginine and lysine (Stable Isotope Labelling of Amino Acids in Culture; SILAC), then analysing as two groups of three variants (Fig 1A, Table S2).

**Figure 1.**
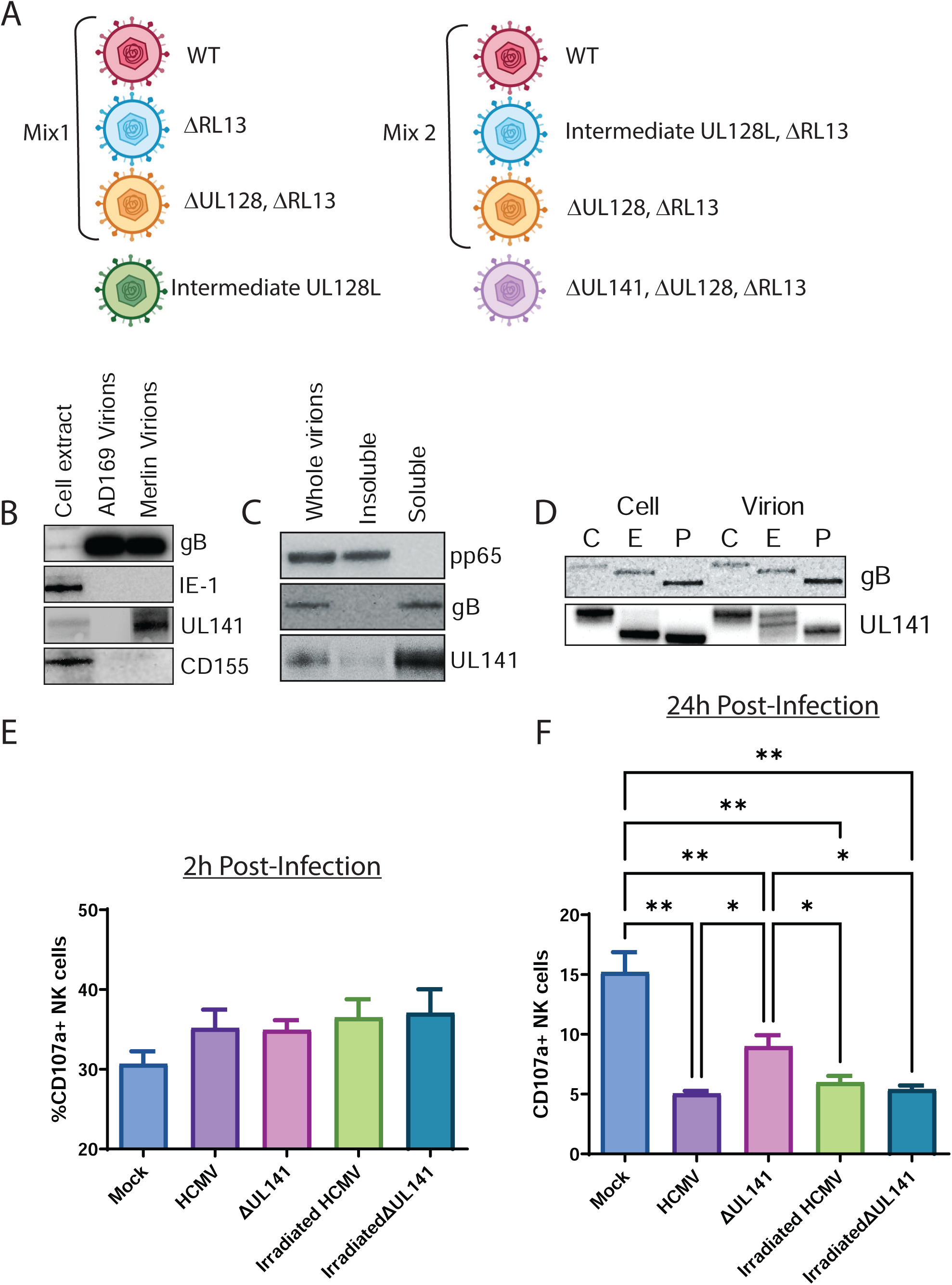
gpUL141’s presence in the virion envelope does not lead to NK inhibition. (A) Schematic of virions analysed in this study. (B) Virions from the passaged AD169 strain, or the wildtype Merlin strain, were purified from HFFF and analysed by Western blot. (C) Virions from strain Merlin were purified from HFFF then treated with detergent to extract membrane proteins, before assessing by Western blot. (D) Virions from strain Merlin were purified from HFFF then left untreated (E) or digested with EndoH (E) or PNGaseF (P) before analysing by Western blot. (E-F) cell-free strain Merlin virions were added to HFFF for 2h (E) or 24h (E), then cells were co-incubated with NK cells for 5h in the presence of CD107a antibody and golgistop, before levels of NK-cell degranulation were assessed.

These SILAC analyses of near identical Merlin virions, differing only in RL13 and UL128L, indicated that deletion of RL13 made very little difference to the relative abundance of virion proteins while loss of UL128L led to higher levels of gO (as previously reported)^32,33^ and UL146 (Figure S1). Aside from this, there were no consistent differences that would support effects of these proteins on other virion components. The absolute Merlin virion proteome was determined as the consensus of proteins identified across all virion preparations. 96 viral proteins were designated as ‘high confidence’ when the peptides were present in at least half of all virus preparations (Table S1). Of these, 53-68 were previously identified by mass spectrometry analyses of AD169 virions^28,29^ and 35 by mass spectrometry analysis of TB40^30,31^. A further four proteins not identified by mass spectrometry analysis of AD169 or TB40 virions, but recognised as HCMV virion components following assessment of virions by alternative approaches, were also detected with high confidence (UL36, gpRL13^8^, gpUL130^34,35^, UL23^36^). 18 proteins identified with high confidence have not been previously described as virion components, with at least 10 membrane-localised. Some of these have been found as virion components in CMVs from other species, however others are unique to HCMV (Table S1).

We used iBAQ (intensity-based absolute quantitation) values for virions with a wild-type genome to estimate the relative abundance of different virion components. Certain proteins accounted for a substantial overall proportion of the total protein content of the virion. These included tegument proteins pp65 (∼43%), pp71 (∼4%) and pp150 (∼1.5%) as well as capsid proteins UL86 (∼5%) and UL85 (∼4%). To determine which viral components were specifically enriched in the virion in contrast to lysates of infected cells, we compared iBAQ values for virion proteins with those we previously calculated for cells infected with Merlin-strain HCMV^22^. Most proteins exhibited similar abundance, but there was a 2-340 fold enrichment of multiple virion membrane proteins, as well as capsid and tegument proteins. These included membrane proteins gN, gM, gH and gL, capsid proteins UL46, UL47 and UL85 as well as tegument proteins UL83, UL82 and UL103 (Fig. S2A-B, Table S3A). There was also enrichment of certain types of cellular proteins, particularly those with transmembrane domains, lipoproteins, ATP synthase subunits and host receptors for viral entry. The latter included known receptors for HCMV including Integrins β1 and α2 (Figure S3A-B, Tables S2B-C).

### gpUL141 is a novel virion envelope glycoprotein, but does not act as an immune-evasin when delivered by the virion

We have previously characterised the viral protein gpUL141 as a potent inhibitor of NK activation. When expressed within the cell, it binds CD155 and all four TRAIL receptors and prevents surface expression by retaining them within the ER^12,14^. This has the effect of rendering cells more resistant to TRAIL-mediated apoptosis and NK-cell attack. It also co-operates with US2 to target CD112 for degradation, preventing NK-cell activation^13,17^. We were therefore interested to note that our proteomics analysis identified gpUL141 as a novel virion component. It may not have previously been identified as such because many HCMV strains have acquired mutations during *in vitro* passage that abrogate UL141 expression^6,37^. Western blot analysis confirmed the presence of gpUL141 in purified virions in the absence of its cellular ligand CD155 (Fig 1B). Fractionation of virions confirmed that it was in the envelope compartment (Fig 1C), consistent with the fact that it encodes a transmembrane domain and a signal peptide. Interestingly, gpUL141 found in the virion was resistant to EndoH, while cell-associated gpUL141 was sensitive (Fig 1D), suggesting that gpUL141 is present as two fractions – one is retained within the ER within the cell, while the other must traffic through the ER and the Golgi to be incorporated into the virion.

Given the characterised role of gpUL141 as an immune-evasin, we investigated whether virion-delivered gpUL141 could act as a rapid immune-evasin, prior to *de novo* gene expression. Cells were infected with wildtype HCMV or HCMV lacking UL141, either as live or irradiated virus. Cells were then co-cultured with NK cells and NK-degranulation assessed. NK activation was not affected by the presence or absence of UL141, nor the ability of the virus to initiate *de novo* gene expression, at 2h post-infection (Fig 1E). At 24h post-infection, loss of UL141 led to increased NK activation, but this was not seen in the irradiated virus (Fig 1F). Thus, this effect was due to gpUL141 synthesized de novo within the infected cell. We therefore concluded that virion delivered gpUL141 does not impact NK cell activity, only gpUL141 expressed *de novo* within the cell exhibited this property.

### gpUL141 interacts with multiple HCMV entry glycoproteins within the virion

To investigate what roles virion-associated gpUL141 might play, we carried out a SILAC-immunoprecipitation (SILAC-IP) in which gpUL141 was pulled down via a C-terminal V5 tag from purified virions, then analysed by MS to identify interacting proteins. A virus lacking pentamer expression was used due to the higher cell-free titres it produces, with analysis revealing interactions of gpUL141 with gB, gH, gL, gO, and gpUL116 (Fig 2A, Table S4). gB is the conserved herpesvirus fusogen, gH/gL/gO form the ‘Trimer’ complex which is essential for cell-free entry into all cell types via interactions with PDGFRa as well as other (unidentified) receptors^38,39^, while gpUL116 interacts with gH but has no known receptor binding role currently^40–42^. All interactions were confirmed by IP-western of gpUL141 from purified virions (Fig 2B). Thus, gpUL141 interacts with multiple members of the HCMV entry machinery within the virion.

**Figure 2.**
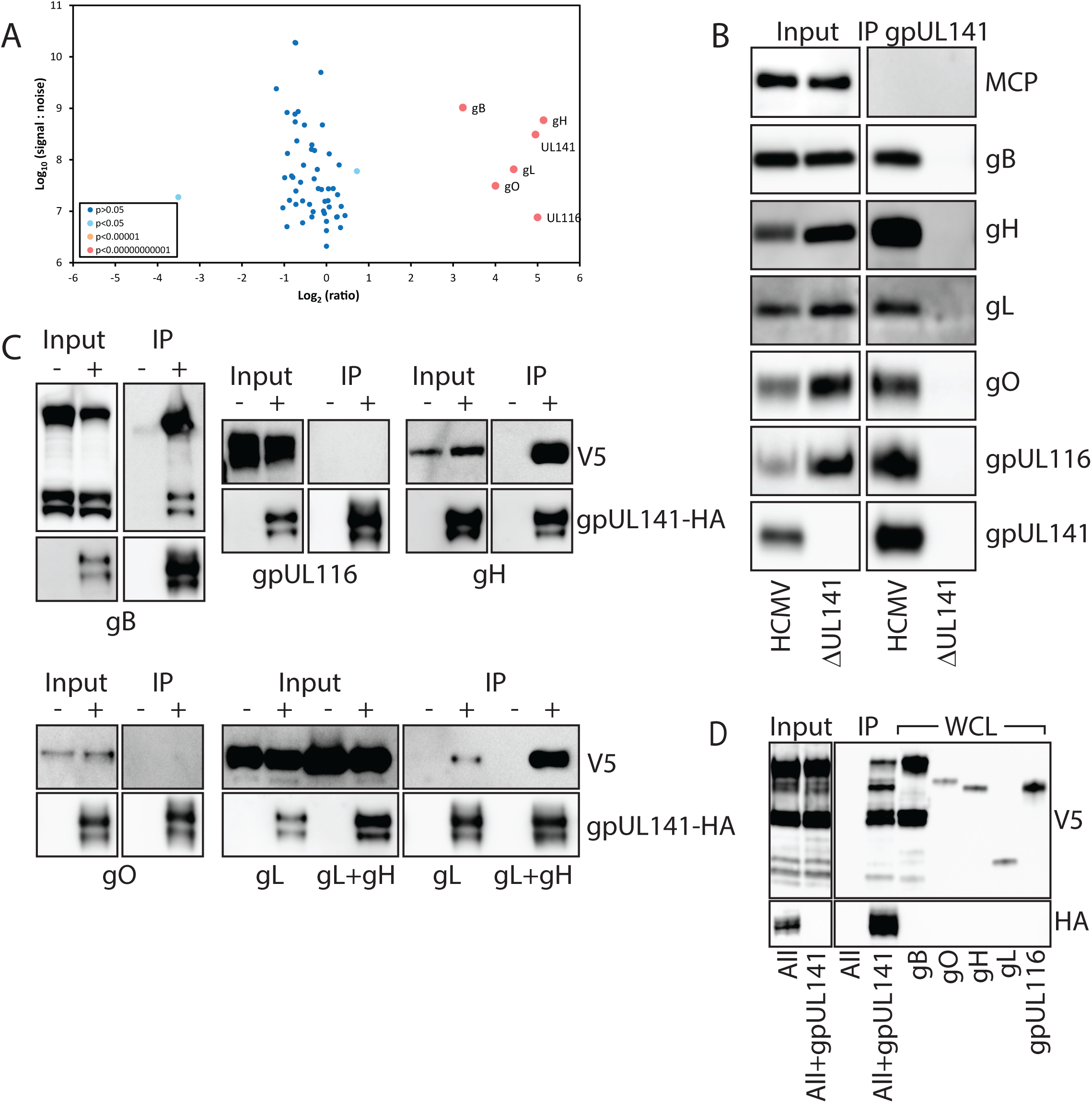
gpUL141 interacts with multiple entry glycoprotein complexes. (A) Cell-free strain Merlin virions that lacked UL128L but contained either a C-terminal V5 tag on gpUL141, or a deletion of UL141, were grown in HFFF in ‘Medium’ or ‘Heavy’ SILAC Media respectively, then purified on glycerol-tartrate gradients and proteins subjected to IP with a V5 antibody. Samples were then analysed by MS, and the ratio of identified proteins between the two samples calculated. (B) Virions were prepared in the same way as (A) but without SILAC media, then analysed by Western blot. (C-D) RAd vectors expressing all potential gpUL141 interactors each with a C-terminal V5 tag, or gpUL141 with a C-terminal HA tag, were used to co-infect HFFF. (C) Each potential interactor was expressed alone (-) or with (+) gpUL141, then gpUL141 IP’d using an anti-HA antibody, before assessing by Western blot. (D) All interacting proteins were expressed together from RAds in the absence or presence of gpUL141, then IP performed for gpUL141, followed by Western blot analysis. To identify individual proteins in the mix based on size, whole-cell-lysates (WCL) of each protein expressed alone are provided alongside.

To determine which interactions with gpUL141 might be direct, gpUL141 was expressed from an adenovirus vector along with each protein from the SILAC-IP individually, then immuno-precipitated. Only gH and gB were robustly co-immunoprecipitated with UL141 in this pairwise test, indicating a potentially direct interaction. Although some interaction was observed with gL, this was significantly weaker than when gH and gL were co-expressed, indicating only a weak interaction. All other proteins are presumably pulled down by indirect interactions via gH/gB (Fig 2C). Consistent with this, when all proteins were expressed together from an adenovirus vector, all co-immunoprecipitated with gpUL141, indicating that no other viral proteins are required for all complexes to form (Fig 2D).

Deep learning structure prediction using AlphaFold3 (AF3)^43^ was used to further understand the potential direct physical interactions between gpUL141 and gH or gB. Previous structural characterisation of gpUL141 has indicated that it forms a head-to-tail homodimer when in complex with its cellular targets CD155 or TRAILR2^44,45^, hence two copies of the gpUL141 sequence were included in all structure predictions. AF3 predicted a high confidence interaction between gH and the gpUL141 dimer, with gpUL141 contacting domains DI, DII and DIII of gH (Fig 3A, Fig S4). This prediction is supported by a recently deposited experimental structure which shows a high level of overlap (Fig S4)^46^. AF3 did not predict a high confidence interaction between gpUL141 and gB (Fig. S4), potentially because AF3 predictions yielded a post-fusion conformation of gB rather than the pre-fusion complex likely to predominate in HCMV virions.

**Figure 3.**
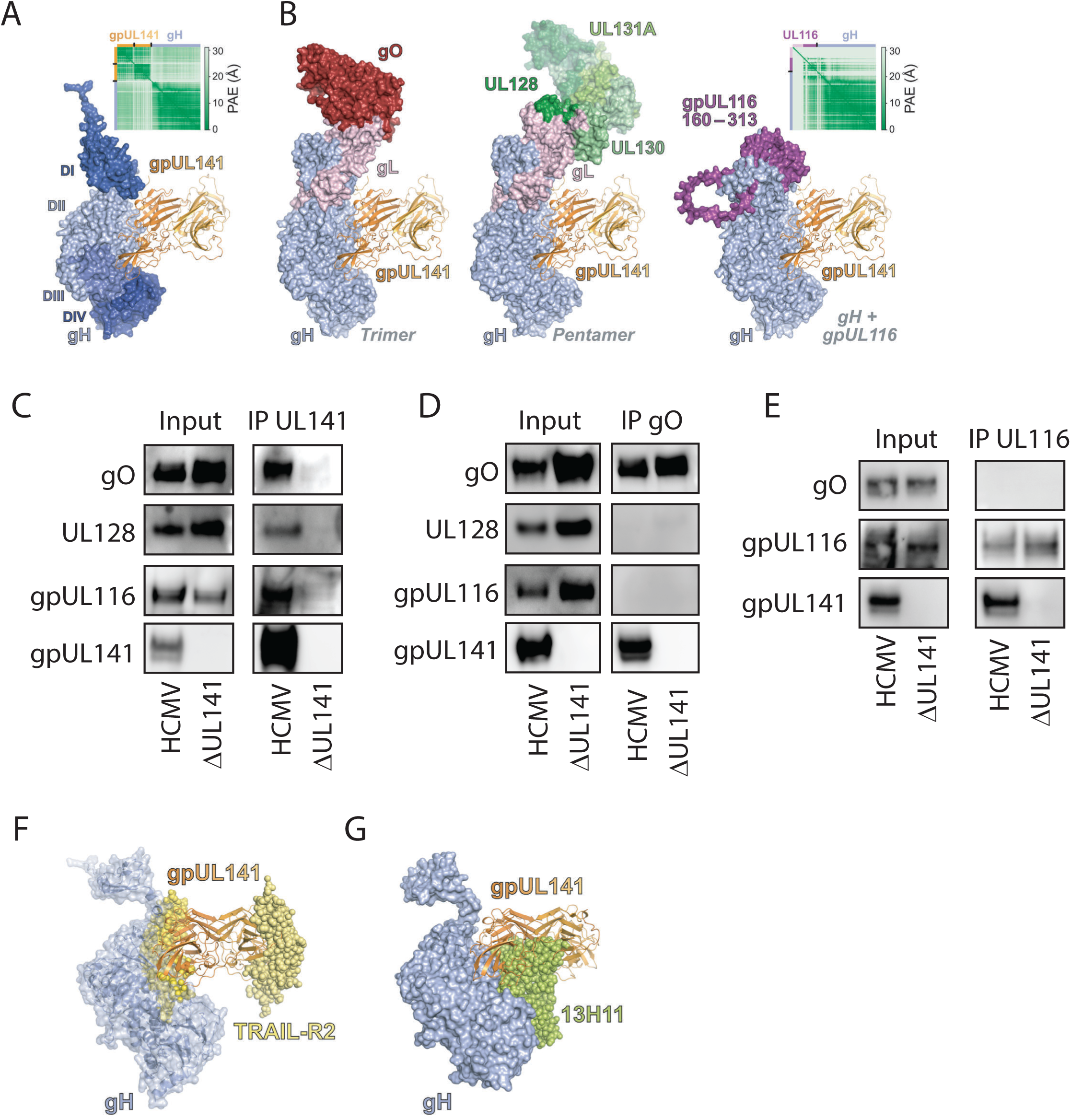
gpUL141 interacts independently with all major gH-containing glycoprotein complexes. (A) Predicted structure of gH (blue surface, coloured by domain) in complex with a homodimer of gpUL141 (orange ribbons). Inset shows the predicted aligned error (PAE) of the model. (B) Superposition of gpUL141 from the predicted gH:gpUL141 structure onto the electron cryo-microscopy structure of the trimer (PDB 7LBE^39^, *left*), the crystal structure of the pentamer (PDB 5VOB^82^, *middle*), and a predicted complex of gH with UL116 (*right*). For clarity, the N-terminal region of UL116 that is predicted to be highly unstructured is not shown. Inset shows the PAE of the gH:UL116 model. (C) HFFF were infected with strain Merlin containing either a C-terminal V5 tag on UL141, or a deletion of UL141, for 96h then proteins subjected to IP with an anti-V5 antibody followed by western blot. (D) HFFF were infected with strain Merlin containing or lacking UL141 expression for 96h, then proteins subjected to IP with an anti-gO antibody followed by western blot. (E) HFFF were infected with strain Merlin expressing a C-terminal HA tag on gpUL116 and either containing or lacking gpUL141 expression for 96h, then proteins subjected to IP with an anti-HA antibody followed by western blot. (F) Superposition of the predicted gH (blue semi-transparent surface and ribbons) plus gpUL141 (orange ribbons) complex onto the crystal structure of gpUL141 in complex with the death receptor TRAIL-R2 (yellow spheres; PDB 4I9X^12^). (G) Superposition of gpUL141 from the predicted gH:gpUL141 structure (orange ribbons) onto the electron cryo-microscopy structure of gH (blue surface) bound to the Fab region of antibody 13H11 (lime green spheres; PDB 7LBE^39^)

### gpUL141 interacts independently with all major gH-containing glycoprotein complexes

HCMV encodes three gH-containing complexes – trimer (gH/gL/gO), pentamer (gH/gL/UL128/UL130/UL131A), and UL116 (gH/UL116) – which are mutually exclusive; UL128 and gO compete for the same binding sites on gL^47^, while gL competes with gpUL116 for binding to gH^40^. The AF3-predicted binding of gpUL141 would not be expected to inhibit the interaction between gH and either gL or gpUL116 (Fig 3B, Fig S4), consistent with our immunoprecipitation experiments, implying that gpUL141 interacts with both gpUL116 and the trimer via gH. This structural model would also imply that gpUL141 could in addition interact with pentamer via gH (Fig 3B). Indeed, when gpUL141 was immunoprecipitated from infected cells, it pulled down gpUL116, gO, and UL128, indicative of interactions with all recognised gH containing complexes (Fig 3C).

The predicted gpUL141:gH structure is consistent with dimers of gpUL141 symmetrically binding two gH molecules. In this case it could potentially ‘bridge’ different gH complexes (Fig. S4). To determine whether this occurs we immunoprecipitated gO, then stained for gpUL116 or UL128. Whereas immunoprecipitation of UL141 pulls down all of these proteins (Fig 2B/3C), pulling down gO did not pull down gpUL116 or UL128 (Fig 3D). Similarly, pulling down gpUL116 did not co-precipitate gO (Fig 3E). Therefore, gpUL141 interacts with all three gH containing complexes (Trimer, Pentamer, and gpUL116), but does not link different complexes together. In contrast, the predicted gH binding interface of gpUL141 overlaps significantly with the surface bound by TRAIL-R2 (Fig 3F), suggesting that binding of gpUL141 to gH (and complexes thereof) and TRAIL death receptors would be mutually exclusive, consistent with virion SILAC-IP data (Fig 2A) and western blot (Fig 1B). Interestingly, the predicted gH/UL141 interface also overlaps with the binding site of a gH neutralising antibody (13H11^48^), indicating that UL141 may impact on epitope availability for antiviral antibodies (Fig 3G).

### gpUL141 modulates cell-surface and virion gH-complex levels

We have previously shown that knocking UL141 out of the virus genome has no apparent impact on cell-free virus levels in fibroblasts, but enhances titres in epithelial cells^37^. This phenomenon was not restricted to epithelial cells (Fig 4A), virus dissemination was also enhanced in the absence of UL141 in endothelial cells (Fig 4B), suggesting that it occurs in all cell-types where entry is dependent on pentamer. Since gpUL141 can inhibit virus dissemination and interacts with multiple gH-containing complexes that are required for virus infection, we investigated whether these phenomena were linked. The majority of gpUL141 is retained within the ER where it prevents transit of CD155 and TRAILR2 through the ER-Golgi apparatus^12,14^, yet our data also indicated that virion envelope glycoproteins must traffic through the ER and Golgi in order to become incorporated in the virion envelope (Fig 4C). We therefore investigated whether the gpUL141 retained in the ER also bound gH-containing complexes and affected their transition through the ER/Golgi. We took advantage of the fact that gO is highly glycosylated, and therefore trafficking from the ER through to the Golgi can be assessed by measuring sensitivity to EndoH. When epithelial cells were infected with HCMV lacking UL141 there was a significant fraction of EndoH resistant gO, which was reduced in virus expressing UL141, indicating that more gO is retained in the ER in the presence of gpUL141 (Fig 4C).

**Figure 4.**
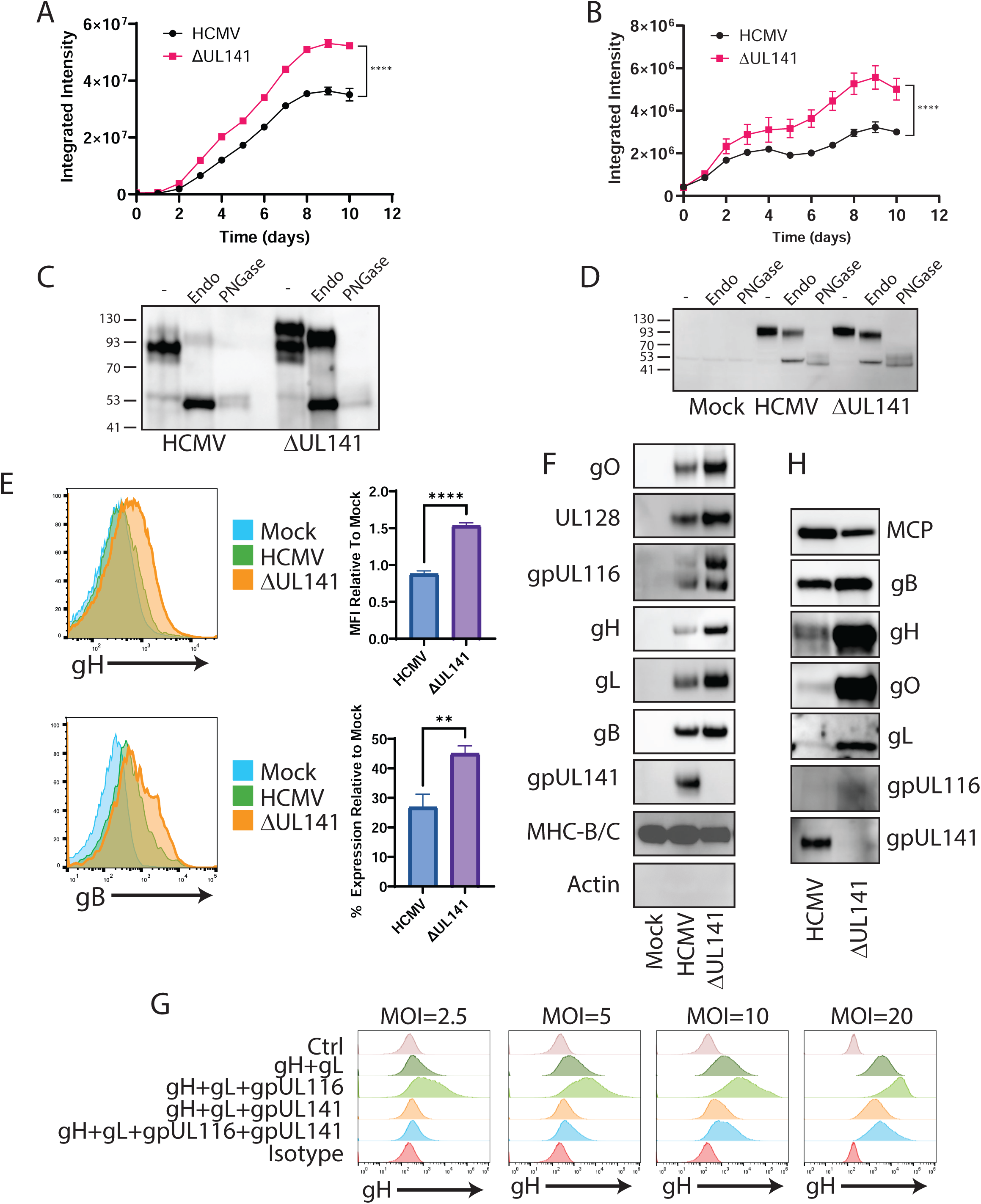
gpUL141 alters the glycoprotein composition of both the virion and the infected cell-surface. (A-B) HFFF were infected with strain Merlin-GFP with UL141 either intact or deleted, at high MOI. 72h later cells were co-cultured with RPE-1 (A) or HUVECs (B). Once the HFFF had lysed (192h), virus dissemination through the RPE-1 or HUVEC monolayer was assessed by incucyte. (C) RPE-1 cells were infected as in (A), 192h post-co-culture infected RPE-1 cells were subjected to digestion with EndoH or PNGaseF, or left under control conditions, then analysed for gO protein by Western blot. (D) HFFF were infected at high MOI with strain Merlin containing or lacking UL141, then 96h later subjected to digestion with EndoH or PNGaseF, or left under control conditions, and analysed for gO protein by Western blot. (E) HFFF were infected for 72h with strain Merlin containing or lacking gpUL141, then levels of gH and gB on the cell surface assessed by flow cytometry. (F) HFFF were infected for 72h at high MOI with strain Merlin containing or lacking UL141, then cell surface proteins isolated by Plasma-membrane profiling (PMP) followed by Western blot. (G) HFFF-CAR were infected with RAd vectors expressing the indicated proteins for 48h, then stained for cell surface gH levels and assessed by flow cytometry. In all cases RAd-UL141 was used at MOI=20, all other RAds were used at the indicated MOI. (H) HFFF were infected for 72h at high MOI with strain Merlin either expressing or lacking gpUL141, then co-cultured with uninfected RPE-1 cells. Virus was allowed to disseminate through the RPE-1 monolayer until 100% CPE was achieved, at which point virions were purified and analysed by Western blot.

Although we had investigated this phenomenon in epithelial cells due to the restriction in virus dissemination observed in this cell type, there was no reason to assume that it was restricted to epithelial cells. When a comparable assay was carried out in fibroblasts, we observed a similar pattern, albeit with a smaller reduction (Fig 4D). We therefore expanded this result by measuring cell-surface levels of gH and gB by flow cytometry in fibroblasts. At 72h post-infection in the presence of gpUL141, gH and gB levels were low (gB) or undetectable (gH). However, when UL141 was deleted, surface levels of gH and gB were enhanced (Fig 4E). Antibodies are not available to detect the cell-surface forms of other gH-containing complexes. To determine whether this phenomenon extended to all proteins that interact either directly or indirectly with gpUL141, we therefore used our previously validated ‘Plasma Membrane Profiling’ technique to isolate cell-surface proteins and assess levels by western blot^16,49^. In support of EndoH and flow analysis, levels of gO, gH, gL, gpUL116, gB, and UL128 on the plasma membrane were all reduced in the presence of gpUL141, while a control protein (free heavy chains of HLA-B/C) was unaffected (Fig 4F). This phenomenon could also be recapitulated outside of the context of infection, using adenovirus vectors expressing individual glycoproteins (Fig 4G). When gH is expressed in isolation it does not traffic to the plasma membrane, but it becomes surface-localised when co-expressed with gL, and this is enhanced when gpUL116 is additionally co-expressed. However, in all cases, co-expression of UL141 resulted in a reduction in cell-surface levels. This was dose-dependent, with an excess of UL141 leading to almost complete inhibition of surface trafficking.

Since cell-surface trafficking was altered in the presence of gpUL141, we wondered whether this would also affect the levels of gH-containing complexes found in purified virions. Indeed, in the input samples used for our virion IP-western (Fig. 2B), the levels of proteins from all complexes were all significantly lower in the presence of gpUL141, with the potential exception of gB. Since the effects of gpUL141 on virus growth were more extensive in epithelial cells, we also tested a subset of proteins in virus derived from this cell type. As in fibroblasts, the levels of proteins that interacted directly or indirectly with gpUL141 were reduced within the virion (Fig 4H). Thus, gpUL141 acts to reduce the incorporation of all gH containing complexes into virions, and limits presentation of these complexes (along with gB), on the plasma membrane of the infected cell.

### gpUL141-mediated modulation of gH-complex levels is due to its ER retention domain

UL141 is retained within the ER where it prevents ER-exit of CD155 and TRAILR^12,14^. It has previously been assumed that this ER-retention function is necessary to prevent cell-surface trafficking of these cellular proteins in order to prevent activation of NK cells. However, our data implies that the ER retention function may have additional impacts on multiple viral entry glycoproteins. For transmembrane proteins, the ER retention domain is often in the C-terminus of the protein, with the classical sequence being KKXX. UL141 contains such a motif at its extreme C-terminus. To explore this further, we therefore generated HCMV viruses in which the C-terminal intracellular portion of UL141 was deleted. We have previously shown that this results in increased trafficking of gpUL141 through the ER/Golgi when expressed from a RAd^10^, however surprisingly it did not inhibit the ability of RAd expressed gpUL141 to reduce cell surface levels of CD155, with levels actually down-regulated to a greater extent when the ER retention domain was deleted (Fig 5A). Similarly, when we engineered this mutation into HCMV, it did not alter the ability of gpUL141 to limit cell-surface levels of CD155, TRAILR2, or CD112 (Fig 5B). Therefore, the ability of UL141 to act as a NK-evasin does not require UL141 to be ER-resident.

**Figure 5.**
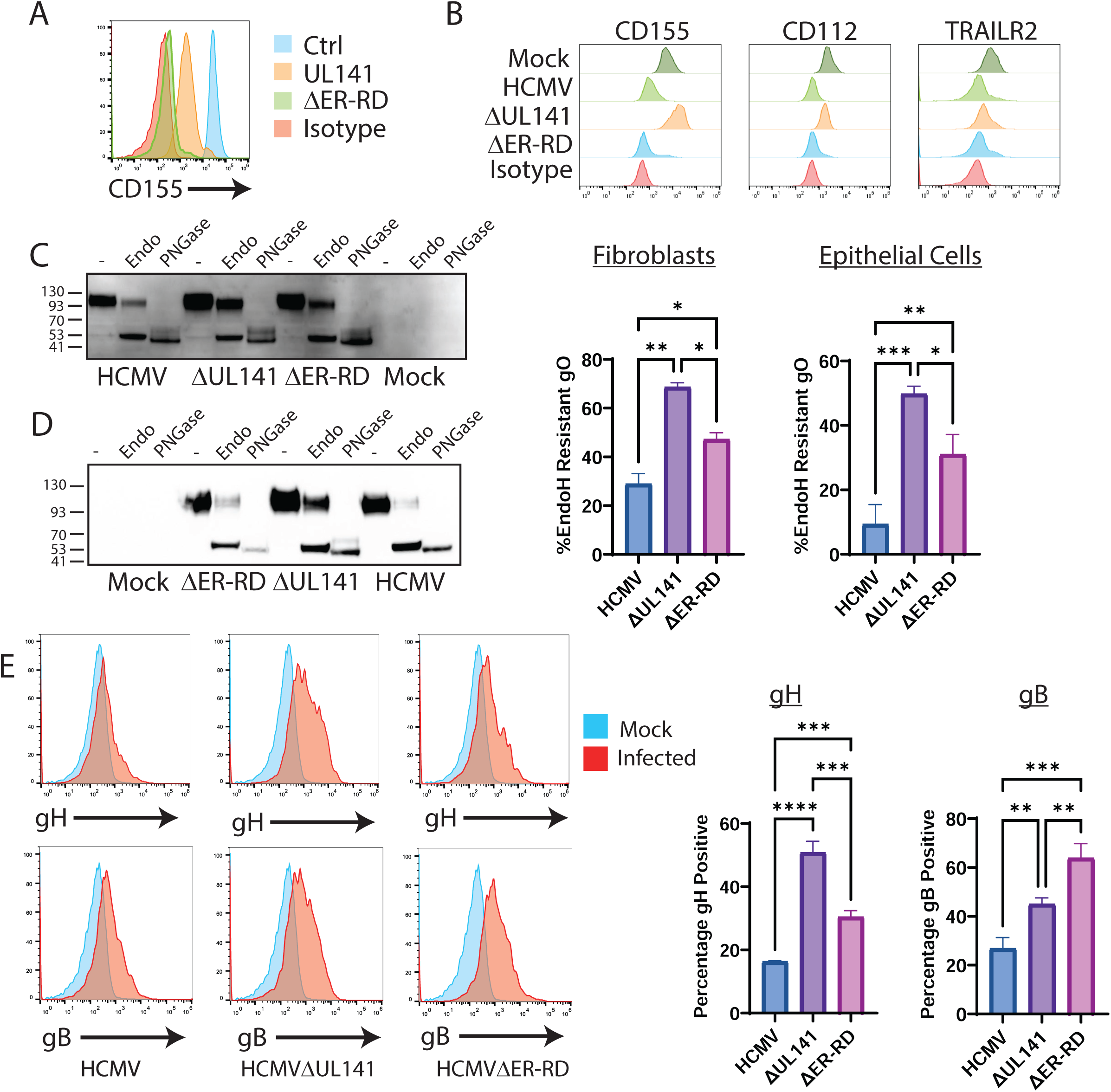
The gpUL141 ER retention domain alters envelope glycoprotein trafficking. (A) HFFF-CAR were infected with RAd vectors expressing wildtype UL141 or UL141 in which the C-terminal intracellular portion (containing an ER-retention domain (ER-RD)) had been deleted, then 48h later were stained for cell-surface CD155 levels and analysed by flow cytometry. (B-F) Cells were infected with wildtype HCMV strain Merlin or virus lacking either the whole of gpUL141, or just the C-terminal intracellular portion of gpUL141 (ER-RD). (A) HFFF were infected with RAd vectors encoding UL141 or ΔER-RD for 48h or (B) with HCMV encoding UL141, deleted for UL141, or ΔER-RD, for 72h and then stained with anti-CD155 (A) or anti-CD155, anti-CD112, or TRAILR2 antibodies (B) and analysed by flow cytometry. (C) HFFF were infected for 96h or (D) HFFF were infected for 72h then co-cultured with uninfected RPE-1 cells for a further 8d, then samples subjected to digestion with EndoH or PNGase F, or left undigested before being analysed by Western blot and stained for gO. (E) HFFF were infected for 96h then stained for cell surface gH or gB followed by flow cytometry.

To determine whether the ER retention domain affected gH-complex trafficking, we repeated the EndoH analysis of gO. In both fibroblast (Fig 5C) and epithelial cells (Fig 5D), loss of the ER retention domain led to an increase in levels of EndoH resistant gO, although not always to the level seen following complete UL141 deletion. When we assessed cell-surface trafficking of gH and gB, removal of the ER retention domain led to recovery of both proteins on the plasma membrane (Fig 5E).

### Intracellular retention of complexes by gpUL141 limits syncytium formation

Viral entry glycoproteins on the infected cell surface can engage with entry receptors on surrounding cells leading to fusion and syncytium formation. In the case of HCMV, this requires gH/gL complexes and gB^50,51^. We reasoned that if all of these were reduced on the infected cell surface, this could reduce receptor engagement and therefore the chances of syncytium formation. ARPE-19 epithelial cells have previously been reported to readily form syncytia upon HCMV infection^51^. These cells were therefore infected with GFP-expressing HCMV, and virus allowed to disseminate. Syncytia were first observed by day 6 post-infection, and well established by 8 days post-infection, at which point a clear difference was seen. Virus expressing UL141 demonstrated significantly lower formation of large multi-nucleated cells in comparison with virus lacking UL141 (Fig 6). Furthermore, the size of syncytia were much lower in wildtype virus as compared to the virus lacking UL141 expression. Deletion of the ER retention domain resulted in syncytia of comparable size and number to ΔUL141 virus, indicating that this phenomenon was due to reduced cell-surface gH-complex trafficking as opposed to the presence of gpUL141 in cell surface complexes. Thus, gpUL141-mediated modulation of entry glycoproteins inhibits syncytium formation.

**Figure 6.**
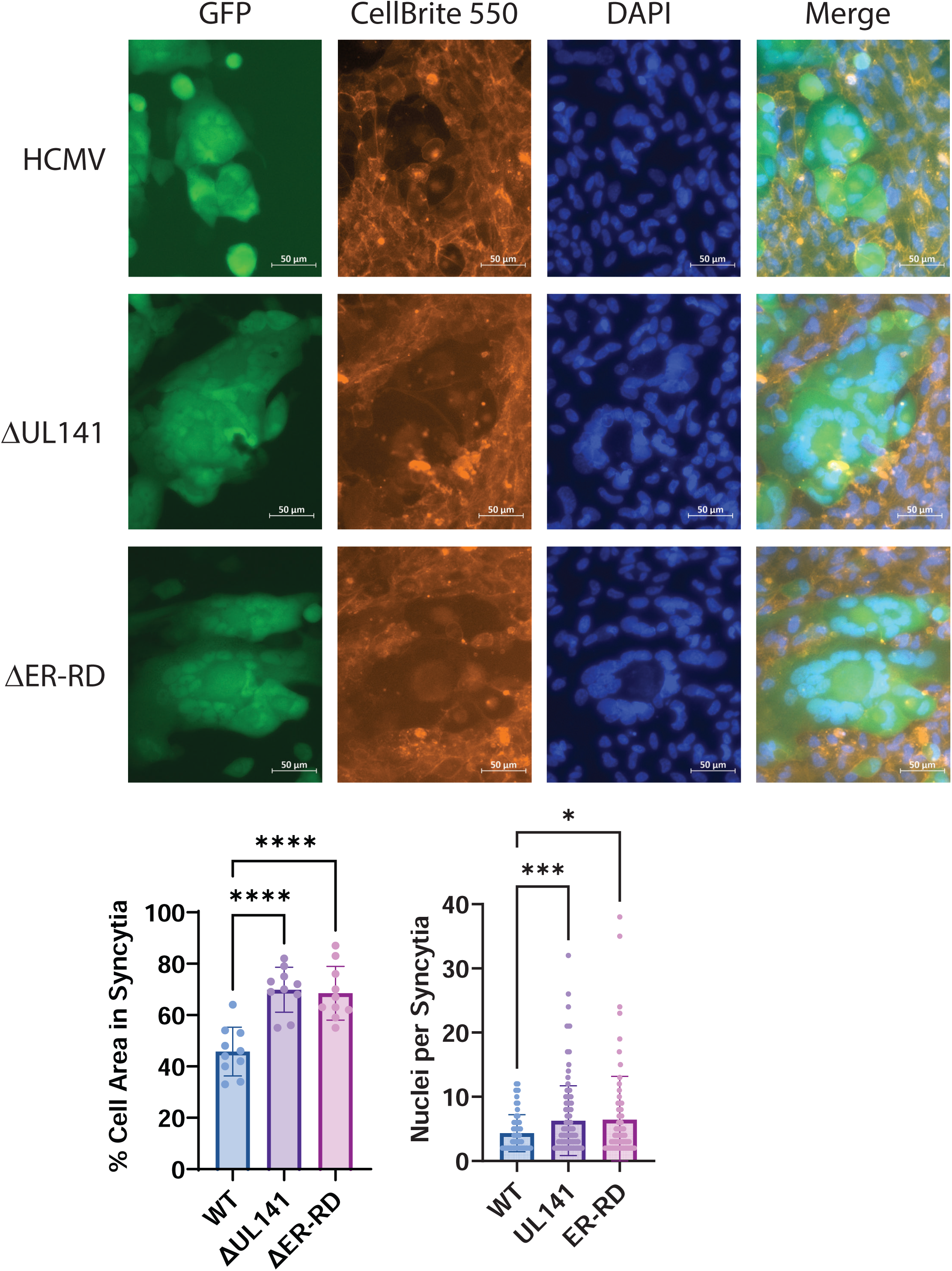
gpUL141 retention of gH complexes limits syncytia formation. HFFF were infected for 72h at high MOI by HCMV strain Merlin expressing GFP and either containing or lacking gpUL141 expression, or expressing an ER retention domain mutant of UL141 (ΔER-RD), then co-cultured with uninfected ARPE19 cells at a ratio of 0.03:1. After a further 8 days nuclei of live cells were stained with Hoechst and CellBrite® 550 Membrane stain, imaged by microscopy, and the number of syncytia quantified.

### gpUL141-mediated modulation of gH-complex levels inhibits humoral antiviral functions

Cell-surface viral antigens are a target for humoral immunity, with opsonised HCMV infected cells susceptible to NK-mediated attack through antibody-dependent cellular cytotoxicity (ADCC). Antibodies capable of mediating ADCC against gB or gH-containing complexes have not been described, therefore to determine whether greater trafficking of antigens to the cell surface can enhance susceptibility to ADCC, we tested levels of ADCC induced by antibodies that we have previously isolated against gpUL141 itself. ADCC was significantly enhanced by the greater cell-surface trafficking that arose from loss of the ER retention domain on UL141 (Fig 7A).

**Figure 7.**
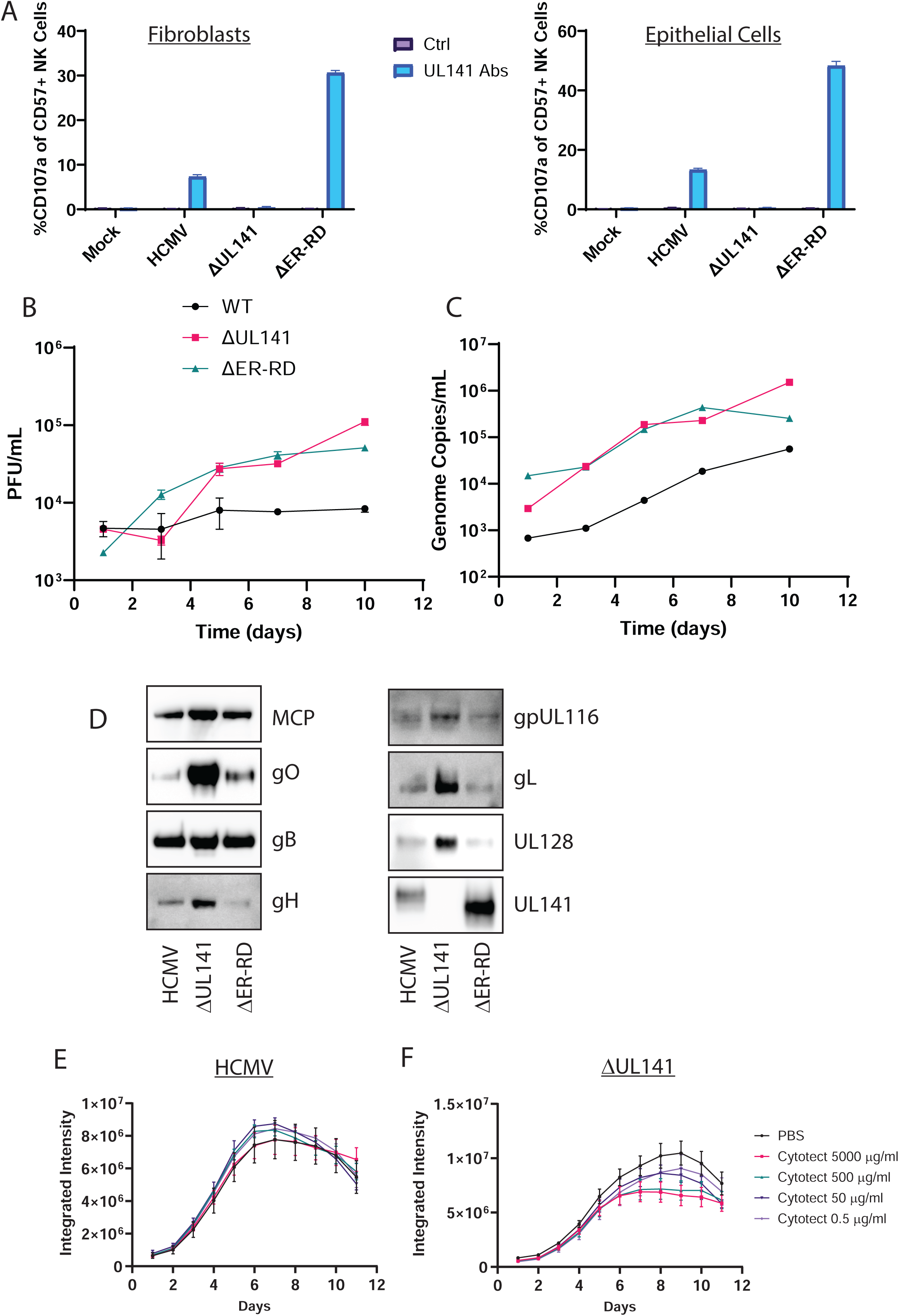
gpUL141 retention of gH-complexes alters virus growth, virion incorporation, and susceptibility to neutralising antibodies. (A) HFFF were infected for 96h, alternatively HFFF were infected for 72h and then co-cultured with RPE-1 cells for 24h before infected RPE-1 cells were isolated by MACs, and left until 96h post co-culture. Infected cells were then co-cultured with NK cells for 5h in the presence of CD107a antibody and golgistop, before the percentage of degranulating NK cells (i.e. those demonstrating ADCC) were assessed. (B-C) HFFF were infected for 72h at high MOI with wildtype HCMV strain Merlin-GFP or virus lacking UL141 then co-cultured with uninfected RPE-1 cells. 5d later the supernatants were harvested and analysed for live virus by plaque assay (B) or encapsidated genomes by qPCR (C). (D) HFFF were infected for 72h with wildtype HCMV strain Merlin or virus lacking either the whole of gpUL141, or just the C-terminal intracellular portion of gpUL141, which contains a ER-retention domain (ER-RD). HFFF were co-cultured with RPE-1 cells and maintained until 100% CPE, at which point cell-free virions were purified and analysed by Western blot. (E-F) HFFF were infected for 72h with HCMV strain Merlin-GFP or the same virus lacking gpUL141, then added to uninfected HUVECs. 5d later the indicated concentrations of cytotect were added, and virus dissemination over time assessed by incucyte.

To determine whether the ER retention domain also altered the growth properties of virus, we assessed cell-free virus titres in epithelial cells. As previously, complete loss of UL141 led to higher cell-free titres, and this was phenocopied by the loss of just the ER retention domain (Fig 7B). This was due to a physical loss of virion secretion because qPCR of viral genomes from the supernatant also demonstrated reductions (Fig 7C). Surprisingly, when the genome:PFU ratio at day 7 was calculated, values of 2.4, 7.2 and 10.6 for WT, ΔUL141 and ΔER-RD respectively, indicate that although UL141 reduced particle release, particle infectivity may be slightly increased. Purification of virions demonstrated that, as in earlier data (Fig 3H), loss of gpUL141 led to increases in levels of all gpUL141-interacting proteins, with loss of just the ER retention domain resulting in increased levels of some gH-complex containing proteins, with gO in particular restored (Fig 7D). Thus, the alteration in growth properties associated with gpUL141 expression arises from the presence of the ER retention domain, which also modulates some virion glycoprotein levels in the virion.

We have previously found that modulating the expression levels of gH-containing complexes alters the sensitivity of HCMV to neutralising antibodies during spread within a monolayer^15^. To determine whether the changes seen here had a similar impact, we infected cells at low MOI with GFP-expressing HCMV strains that either contained or lacked UL141, and measured dissemination over time in the presence of Cytotect, a pooled polyclonal preparation of antibodies from HCMV seropositive individuals and selected for high neutralising activity. Whereas wildtype virus was completely resistant to neutralising antibodies, virus lacking UL141 was susceptible (Fig 7E).

Therefore, gpUL141 contains an ER retention domain which is not required for it to modulate ligands for activating receptors on NK cells, but acts to restrict cell-surface expression of gB and all gH-containing glycoprotein complexes, while also affecting virion envelope composition. This reduces virus dissemination, but renders it more resistant to both neutralising antibodies and ADCC. Thus, UL141 acts as an evasin of both innate (NK) and adaptive (humoral) immunity.

## Discussion

Our analysis significantly expands the repertoire of proteins found in the HCMV virion, adding at least 18 viral proteins found in the majority of highly purified preparations, at equal or greater abundance than established components. Not all detected proteins need be packaged specifically in the virion, cellular RNAs for example can be packaged in virions in proportion to their relative abundance, indicating a non-specific process^52^. Nevertheless, a significant number of cellular proteins were enriched in the virion, implying their specific recruitment either to the virion itself or to sites of virion assembly. Furthermore, gpUL141 provides an exemplar of the novel viral proteins identified, demonstrating that they can be *bona fide* components that play key functional roles in virion assembly and function.

gpUL141 was previously recognised to promote virus survival through three distinct mechanisms (targeting CD155^14^, CD112^13^, all four TRAIL receptors^12^). As cell and virion surface proteins, there are multiple examples of viral entry glycoproteins being ‘co-opted’ to also act as immune evasins^53^. However, although gpUL141 is found in the virion in complex with nearly all HCMV entry glycoprotein complexes, our data argues that rather than acting in entry, gpUL141 acts to modulate the levels of other glycoproteins in order to control syncytium formation and avoid both neutralising and non-neutralising antibody-mediated control. Given that gpUL141 interacts with multiple host cell-surface proteins, a potential function for virion-presented gpUL141 could involve receptor-mediated viral entry. However, given that the presence of gpUL141 results in reduced virus dissemination and lower cell-free titres in epithelial and endothelial cells *in vitro*, any such functions do not seem to be proviral. Furthermore, the ability of gpUL141 to restrict cell-free virus release was lost when the ER retention domain was removed, implying that this phenotype arises from influences on glycoprotein trafficking rather than receptor binding. It is also notable that the structural model indicates that gpUL141 cannot bind to both viral (gH) and cellular (TRAIL-R2) factors simultaneously. Nevertheless, the presence of gpUL141 in the virion does have the potential to influence virus biology through the dimerization revealed by the structural models. It is notable that although gpUL141 reduced virus particle release and envelope glycoprotein incorporation, it did not reduce per-particle infectiousness. It is possible that dimerization of entry complexes increases avidity to compensate for reductions in abundance. Furthermore, since UL141 occludes at least one neutralising epitope on gH^48^ incorporation into the virion may directly influence epitope-specific immunity.

As a large and complex virus with a broad tropism, HCMV encodes over 10 different envelope glycoproteins that form the core receptor binding and envelope fusion functions^54^. gB transits via the plasma membrane before being internalised to be delivered to the viral assembly complex (VAC) for virion incorporation^55^, and presumably the same is true for other glycoproteins. All are therefore potential targets for ADCC. Through targeting two viral proteins (gH and gB) that are common to nearly all of these complexes, gpUL141 provides an elegant solution to this problem. Both knock-in (RAd) and knock-out (HCMV; both whole protein and ER-RD) support a model in which gpUL141 inhibits plasma-membrane transport and subsequent virion incorporation of the majority of HCMV entry glycoproteins. We have previously shown that the humoral ADCC response during natural infection is highly polyclonal, targeting numerous proteins from the early phase of the viral lifecycle onwards^10^. ADCC activity peaked prior to the expression of entry glycoproteins. This could be in part due to UL141 limiting glycoprotein levels, however it is also possible that UL141-mediated intracellular retention is important to limit the generation of ADCC-antibodies against entry glycoproteins during infection. Dissecting this will require isolating and mapping monoclonal antibodies that target gB and Trimer/Pentamer/gpUL116 that are ADCC-capable, as we have previously for gpUL141. However, given the large number of antigens that gpUL141 influences, even small alterations in cell-surface levels have the potential to produce significant effects.

Although complete loss of gpUL141 resulted in robust effects on plasma membrane and virion envelope composition, the impact of removing just the ER-RD was more varied. Modulation of plasma membrane levels were either partially or entirely dependent on the ER-RD, depending on the assay and protein being tested. This likely reflects the presence of multiple different competing signals within the C-terminal intracellular domain of gpUL141, all of which were lost by complete deletion. In addition to the ER retention dilysine motif in it’s C-terminus, gpUL141 also encodes a YxxΦ motif for endosomal retrieval from the plasma membrane^56^. ER trafficking can also depend on interactions between C-terminal and membrane domains^57^. Thus, gpUL141 may both limit ER-exit of complexes while also promoting re-internalisation into the VAC of those that do traffic to the cell surface. Loss of both motifs may influence multiple processes, with the final outcome also depending on the motifs found in component proteins of the different complexes – for example gB has its own retrieval motif^55^. Assessing this will require more in-depth analysis and targeted mutagenesis of the gpUL141 cytoplasmic tail.

Despite the partial effects seen biochemically following ER-RD deletion, effects on levels of cell-free virus were almost entirely recovered. This may indicate a threshold effect, in which levels are restored sufficiently to improve virus release – in particular gO, which is required for cell-free infection, was upregulated in this virus. These assays imply that for HCMV, reducing envelope glycoprotein levels in the virion may impact virus release, but this is a price worth paying if it also results in lower cell-surface antigenic targets for ADCC, resistance to neutralising antibodies, and lower syncytium formation. The role of syncytia *in vivo* is unclear, however HCMV strains that lack UL141 form syncytia in a range of cell types *in vitro*^58–61^. It is clear from these studies that syncytium formation depends on both cell type and genetic variants of viral proteins including gB^62,63^. Our data imply that syncytia may be detrimental to HCMV *in vivo* since UL141 specifically reduces it. However, although UL141 may modulate the extent of these effects, it is unlikely to completely prevent them since we still saw small syncytia (∼2 nuclei) during infection of ARPE19 cells with wildtype virus.

The levels of multiple envelope glycoprotein complexes are notably different in Merlin compared to passaged HCMV strains. As a ratio, Merlin has higher levels of pentamer and lower levels of trimer^32,33^, and expresses gpRL13^8^. At least some of these differences arise from *in vitro* acquired mutations in passaged strains^8,32^; whether they can also arise from genuine strain-specific effects remains to be determined. Nevertheless, they result in Merlin disseminating via a form of cell-cell spread that is exceptionally resistant to neutralising antibodies^8,15^. Subsequent studies have shown that although these differences explain some of the antibody-resistance of Merlin, they are not the only factor^64^; gpUL141 provides an explanation for these observations. There may also be interplay between gpUL141 and other viral proteins that modulate glycoprotein trafficking. UL148 leads to reduction of pentamer levels in the virion in favour of trimer^65^ by preventing gO degradation^66^, while gpUL116 acts as a chaperone for gH to get to the cell surface^41,42^ and trimer incorporation into the virion^40^. Further work will be needed to investigate these impacts in the presence of UL141.

Our work also explains the long-standing observation that the UL140-145 genome region is frequently lost when HCMV is passaged *in vitro.* Mutations adjacent-to or involving UL141 have been seen in clinical strains passaged in fibroblast, epithelial, and endothelial cells^6^, when Merlin was passaged in epithelial cells^37^, and in multiple laboratory strains including TB40^14,67^, VR1814 and the derived FIX-BAC^6^, and all AD169 variants^68^, many of which were fibroblast-derived. UL141 is clearly detrimental to virus release and dissemination in epithelial and endothelial cells, however these studies imply that it may also be detrimental in fibroblasts. No growth phenotype was associated with UL141 in Merlin in fibroblasts, possibly because the effects on glycoprotein trafficking were weaker than in epithelial cells. Nevertheless, gpUL141 can clearly reduce glycoprotein incorporation into the virion in fibroblasts. gO is hypervariable^69,70^ and gO variants influence other glycoproteins as well as virus growth in diverse ways^33,71–73^. Therefore, different phenotypes may be observed when viruses encode other gO genotypes.

Overall, we have defined the proteome of a wildtype HCMV virion, and demonstrate that novel components encoded in clinical virus have important biological effects. One of these (gpUL141) represents a novel viral strategy for immune evasion which impacts numerous aspects of the lifecycle that are relevant to therapeutic and vaccine design. It is therefore important that genetically intact HCMV strains are tested during preclinical development.

## Methods

### Cell Culture

Human foetal foreskin fibroblasts (HFFFs) immortalised with human telomerase reverse transcriptase (HF-TERTs)^74^, HF-TERT-immortalized HFFFs expressing Coxsackie adenovirus receptor (HF-CAR)^15^, retinal pigment epethilial-1 (RPE-1 and ARPE19) cells were maintained under standard conditions in DMEM (Sigma) supplemented with 10% heat-inactivated fetal bovine serum (Sigma). Human umbilical vein endothelial cells (HUVEC) were maintained under standard conditions in Endothelial Cell Growth Medium (Basal medium plus individual supplements; PromoCell). All human cell lines were deidentified prior to use.

### Viruses and Infections

HCMV variants were recovered from an infectious bacterial artificial chromosome (BAC) clone of strain Merlin. This encodes the complete strain Merlin genome, which was repaired to match the sequence of the original clinical material^8^. Mutations are rapidly selected in the genes encoding RL13 and UL128L upon passage *in vitro*, therefore the promoters for these genes have been rendered responsive to the tetracycline repressor (TetR). In cells expressing tetR, expression is restricted, allowing recovery of genetically intact virus^37^. This virus loses tropism for cell types other than fibroblasts due to repression of UL128L, but can readily infect fibroblasts. If these fibroblasts lack tetR then UL128L is expressed, and virus can be transferred to other cell types (e.g. endothelial and epithelial cells) by co-culture. Deletion of the UL141 ER retention domain was carried out by en-passant mutagenesis^75^ with primers 5’- ACACACTGCATTTTTTAACATCTTATTTTTTTATTTTATGCGTGTTCTCAACAGCACTGCAGGTAACACATAGGGATAACAGGGTAATCGATTT-3’ and 5’- TTTTGGAGTATTTTCACCGTATGTTTCCTATGCTACCTGTGTTACCTGCAGTGCTGTTGAGAACACGCATAAAATAAGCCAGTGTTACAACCAATTAACC-3’. A V5 epitope tag was added to the C-terminus of UL141 by recombineering as previously described^8^, using primers 5’- AGGGGACGACGAGGCGGTGAGGGCTATCGACGCCTACCGACTTACGATAGTTACCCCGGTGTTAAAAAGATGAAGAGGCCTGTGACGGAAGATCACTTCG-3’ and 5’- GCATATTTTAATCACACTATTCACATTTCACACACTGCATTTTTTAACATCTTATTTTTTTATTTTATGCGTGTTCTCACTGAGGTTCTTATGGCTCTTG-3’ to insert the SacB cassette and primers 5’- GACGCCTACCGACTTACGATAGTTACCCCGGTGTTAAAAAGATGAAGAGGGGCTCCGGGGGGTCGGGTGGAAGTGGCGGTAAGCCAATCCCTAACCCGCT-3’ and 5’- ACACACTGCATTTTTTAACATCTTATTTTTTTATTTTATGCGTGTTCTCACGTAGAATCAAGACCTAGGAGCGGGTTAGGGATTGGCTTACCGCCACTTC-3’ to replace the cassette with a V5 epitope tag. A HA tag was added to the C-terminus of UL116 by en-passant mutagenesis using primers 5’- ACCTGAGTGCCAACTTTTGGCGCCAACTGGCTCCTTACCGTCACACTCTCATCGTGCCGCAGACTAGCGCTTACCCCTACGACGTGCCCGACTACGCCTG-3’ and 5’- CAACACCACAGCAGTATCACCGGTCCAGGTGAGAAAGAGAAGCCGCAATCCGGGCGGCGGCACATCAGGCGTAGTCGGGCACGTCGTAGGGGTAAGCGCT-3’. Replication deficient Adenovirus vectors were constructed in the AdZ system as previously described^76^. All modifications were verified by sanger sequencing of the modified site, and all viruses underwent whole genome sequencing following recovery from the BAC^37^.

Infectious HCMV was recovered by transfecting HF-Tet cells with an Amaxa Nucleofector, the basic fibroblast kit, and program T-16. Virus was harvested from the supernatant when 100% CPE was achieved, then cells remove by low speed centrifugation (420 x*g*, 3 min), before virus was pelleted by high speed centrifugation (29,500 x*g*, 2h). Virus was resuspended in DMEM containing 10% FCS, aliquoted, and stored at -80°C. Cell-free titres were determined using an IE1 micro plaque assay. Briefly, HF-Terts were seeded in 96-well plates 18 h prior to use. Virus supernatants were 10-fold serially diluted in DMEM-FBS and added to cells. At 24 h p.i. media was removed and cells fixed in ice cold 50:50 Acetone/Methanol for 15 min. Cells were washed once in PBS and primary antibody added at 1:1000 in PBS for 30 min at 37°C. Cells were washed once in PBS and secondary antibody added at 1:500 in PBS for 30 min at 37°C, followed by a final wash in PBS. Infected cells were counted using an Incucyte® (see below).

For proteomics and biochemical analysis cell-free virus was harvested from the supernatant, cell debris removed by low-speed centrifugation (420 xg, 3 min), then samples concentrated either by dialysis (Viva flow 50R, Sartorius, 1MDa cut-off) or by pelleting through a 20% sorbitol cushion (82,700 x*g*, 1 h). Intact virions were then separated from cellular debris, dense bodies, and non-infectious particles by positive density/negative viscosity gradient centrifugation, as previously described^77^.

Unless otherwise stated HCMV infection of HF-Terts was at MOI=5 for 72 h. To infect RPE-1 or HUVEC cells, HF-Terts infected for 72h were co-cultured with target cells at ratios of 0.03:1, 1:2, or 4:1 HF-Tert:RPE-1/HUVEC for virus dissemination and syncytia analysis, growth kinetics, or PNGase F assays, respectively. For dissemination inhibition assays in HUVEC cells, Cytotect® was added to cells at 3 days post co-culture in a 10-fold serial dilution starting at 5000 µg/mL. HF-CARs were infected with RAd at MOIs stated, for 48 h.

### Proteomics

For analysis of virions by MS, cells were passaged for two weeks in SILAC-DMEM containing arginine and lysine in either their normal, medium (13C6), or heavy (13C6-15N4, 13C6 15N2 respectively) forms respectively, before being infected and virus harvested and purified as above. Following purification samples were lysed in LDS sample buffer and run 1.5cm into a Bis-Tris Midi Gel (Thermo), before lanes were excised with a clean scalpel and cut into 6 equally sized bands. Samples were reduced, alkylated and digested in-gel using trypsin. Digested peptides were eluted with MeCN/water and 5% FA washes. Peptides were pooled in 0.5ml tubes (Protein LoBind, Eppendorf) and dried almost to completion. Samples were re-suspended in 15 µl solvent (3% MeCN, 0.1% TFA) with 7µl analysed by LC-MSMS using a Thermo Q Exactive mass spectrometer (Thermo Fischer Scientific) equipped with an EASYspray source and coupled to an RSLC3000 nano UPLC (Thermo Fischer Scientific). Peptides were fractionated using a 50cm C18 PepMap EASYspray column maintained at 35°C with a solvent flow rate of 250nl/min. A gradient was formed using solvent A (0.1% formic acid) and solvent B (80% acetonitrile, 0.1% formic acid) rising from 3% to 40% solvent B by 90 min followed by a 4 min wash at 95% solvent B. MS spectra were acquired at 70,000 resolution between m/z 400 and 1650 with MSMS spectra acquired at 17,500 fwhm following HCD activation. Data was processed in Maxquant 2.4.8.0 with carbamidomethyl (C) set as a fixed modification and oxidation (M) and acetyl (protein N-terminus) set as variable modifications. Data was searched against a Uniprot Homo Sapien database (downloaded 27/05/14) a curated database of canonical HCMV proteins and a database of common contaminants.

### Separation of Subpopulations Following Co-culture

Where it was necessary to separate cell types following virus transfer by co-culture (e.g. for CD107a assays), a previously established magnetic-activated cell sorting (MACS) separation protocol was used^78^. In brief, HFFF expressing His-tagged mCherry protein on their cell surface were infected with HCMV for 72h, then co-cultured with target cells. Twenty-four hours post co-culture cell monolayers were washed and detached, stained with anti-His antibody (Antibody and Reagents Table) followed by anti-mouse-IgG magnetic beads (Miltenyi Biotec), and separated by MACS (Miltenyi Biotec). Newly infected RPE-1 or HUVEC cells were maintained for a further 48 h prior to CD107a assay.

### Immunoprecipitation

Cells or viruses were lysed in IP Lysis Buffer (Pierce) for 10 min in the presence of protease inhibitors (Sigma), then nuclei removed by centrifugation. Lysates were incubated with anti-V5 or anti-HA agarose beads (Abcam) overnight, before being washed 3 times in lysis buffer and proteins eluted by boiling for 10 min in 4X LDS sample buffer (Thermo Fisher).

### PNGaseF and EndoH digestion

Samples were digested according to manufacturers instructions (NEB). In brief, samples were lysed in denaturing buffer then the appropriate reaction buffer added, and incubated overnight at 37°C in the presence of enzyme. Samples without enzyme added served as controls. The next day LDS sample buffer was added and samples boiled for 10 min before being analysed by SDS-PAGE and western blot.

### Western blot

Samples were lysed in 4X LDS sample buffer (Thermo Fisher) then boiled for 10 min before being loaded onto Bis-Tris Midi gels (Thermo Fisher) and run for 1 h at 200V. Following separation samples were transferred onto PVDF (Amersham) by semi-dry blotting, then blocked in 5% Milk/PBST (blocking buffer) for 1 h. Primary antibodies were added in blocking buffer for 1 h at room temperature or overnight at 4°C, washed 3x in PBST, then secondary HRP-conjugated antibodies added in blocking buffer for 1 h. After washing 3x in PBST membranes were incubated with supersignal west pico (Thermo Fisher) and imaged on a Syngene XX6 geldoc.

### CD107a Assay

PBMCs were isolated from healthy donors and rested overnight in RPMI supplemented with 10% FCS, and L-glutamine (2 mM). Target cells were harvested using TrypLE Express (Thermo Fisher), then mixed with PBMC at an effector:target ratio of 10:1 in the presence of golgistop (0.7 μl/ml, BD Biosciences) anti-CD107a–FITC (clone H4A3, BioLegend), and relevant anti-HCMV antibodies. Cells were incubated for 5 h at 37°C, washed in cold PBS, and stained with live/dead Fixable Aqua (Thermo Fisher), anti-CD3–PECy7 (clone UCHT1, BioLegend), anti-CD56–BV605 (clone 5.1H11, BioLegend), anti-CD57–APC (clone HNK-1, BioLegend). Data were acquired using an Attune NxT Flow Cytometer (Thermo Fisher) and analysed with FlowJo software version 10 (FlowJo LLC).

### Flow Cytometry

Cells were washed in PBS, dissociated with TrypLE Express (Thermo Fisher) and stained with primary antibody (Antibody and Reagents Table) for 30 min at 4°C. Cells were washed in PBS and incubated with the secondary antibody for 30 min at 4°C before being washed and fixed with 4% paraformaldehyde (PFA). All data was acquired using an Attune Nxt Flow Cytometer (Thermo Fisher), and analysed using FlowJo Software version 10 (FlowJo LLC).

### qPCR

Viral DNA was extracted from DNase I treated (RQ1 RNase-free DNase; Promega) supernatants with the QIAamp MinElute Virus Spin Kit (Qiagen) as per manufacturers’ instructions. Quantitative PCR reactions were performed in a Applied Biosystems Quant Studio 3 Thermal Cycler (Thermo Fisher Scientific). Reactions were set up using FastGene 2x IC Green qPCR Universal Mix (Nippon Genetics; Geneflow), 0.4 µM of forward and reverse primers, and 100 ng of cDNA in a final volume of 20 μL. The gB forward primer was 5′**-**CTGCGTGATATGAACGTGAAGG-3′ and the reverse primer was 5′**-**ACTGCACGTACGAGCTGTTGG-3′. Amplification was performed using the Comparative cT with Melt programme (2 min at 50°C, 10 min at 95°C, followed by 40 cycles of 15 sec at 95°C and 1 min at 60°C. Melt curve: 15 sec at 95°C, 1 min at 60°C and 15 sec at 95°C). Water-only negative controls and a serial dilution of a positive control standard (plasmid containing gB) were included in each run. CMV genome equivalents were calculated from the Ct values generated in The Standard Curve App (Thermo Fisher Scientific Connect Data Analysis Apps).

### Incucyte®

Virus dissemination assays and virus titrations were measured on an Incucyte® SX5 Live Cell Analysis System, scanning the whole well with a 4X lens, with both brightfield and green image channels. Wells were scanned every 24 h for 10 days for dissemination and neutralisation assays, or one-off scans for titrations. Incucyte® Basic Analysis Software was used to measure the Total Integrated Intensity (GCU x µ^2^/well) for time course analyses, or Count (per well) for titrations.

### Syncytia Imaging and Analysis

Images were taken from 10 randomly selected fields of view at 20X magnification on a Zeiss Apotome 2 microscope. Syncytia were determined by the presence of two or more nuclei observable within a single cell membrane. Cell area was measured using Zeiss Zen Lite v3.10 software and nuclei counted manually. The average area of fluorescence within syncytia was expressed as a percentage of the total fluorescent cell area.

### Structure prediction

The structures of gH (UniProt Q6SW67 residues 25–719) in complex with gpUL141 (UniProt Q6RJQ3 residues 37–278) and gpUL116 (UniProt Q6SW34 residues 25–313), or of a trimer of gB (UniProt F5HB53 residues 25–751) in complex were predicted using AlphaFold3^43^. For all, predicted signal sequences and transmembrane regions were excluded from the sequence used for structure prediction. Structures were superposed and molecular images were generated using PyMOL (Schrödinger) or COOT^79^.

### Data availability

Atomic coordinates and per-residue quality statistics for the AlphaFold3 models have been deposited in the University of Cambridge Apollo repository (DOI: 10.17863/CAM.118380). Raw proteomics files are available at PRiDE (DOI: PXD065223).

### Statistics

All statistical analyses were carried out in GraphPad Prism v10.4.0. All assays were carried out in a minimum of technical triplicate, and repeated at least 3 times. Statistical tests carried out for each assay are noted in figure legends.

### Antibody and Reagents Table

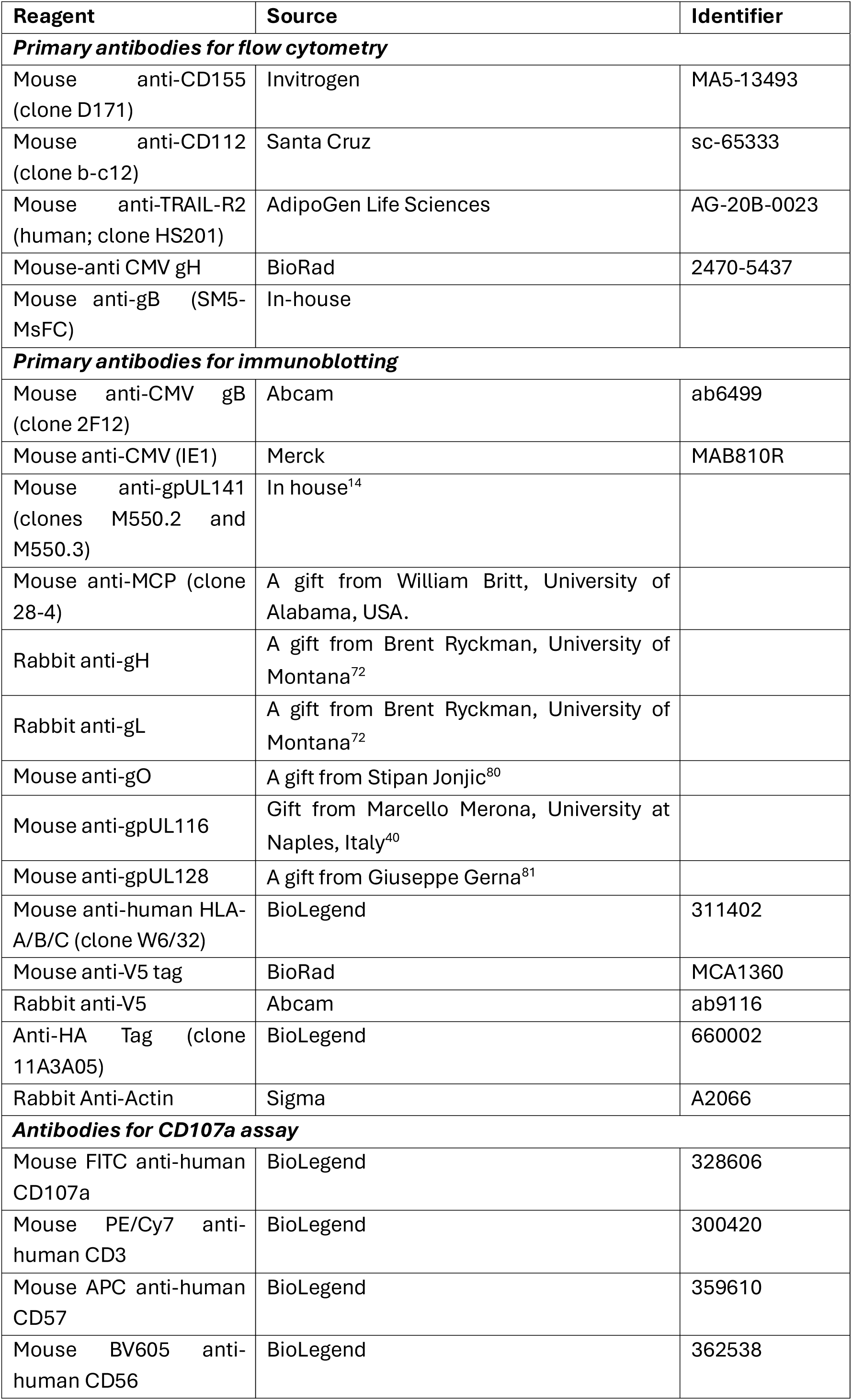
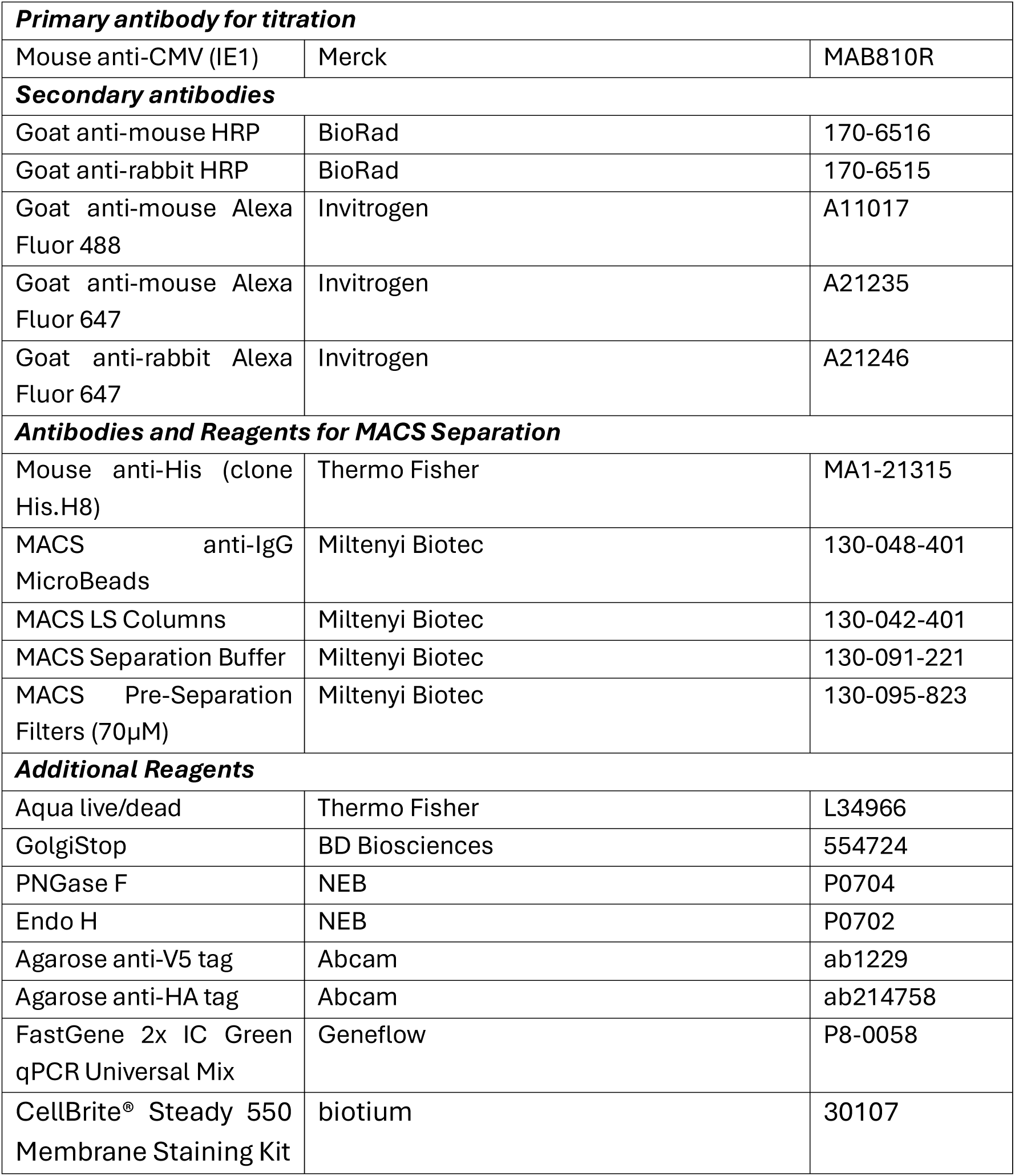

## Supplementary Tables

Table S1. Virus encoded proteins identified by MS-analysis of Merlin virions

Table S2. Mass spectrometry analysis of Merlin virions.

Table S3. (A) HCMV proteins detected in virion or WCL preparations, or both. (B) Comparison of abundance of human proteins detected in virion and WCL preparations. (C) Significantly enriched clusters and their components from DAVID analysis of human proteins enriched with a virion : WCL ratio of >2.

Table S4. Mass spectrometry analysis of SILAC IP of gpUL141 from Merlin Virions.

## Supplementary Figures

**Figure S1:**
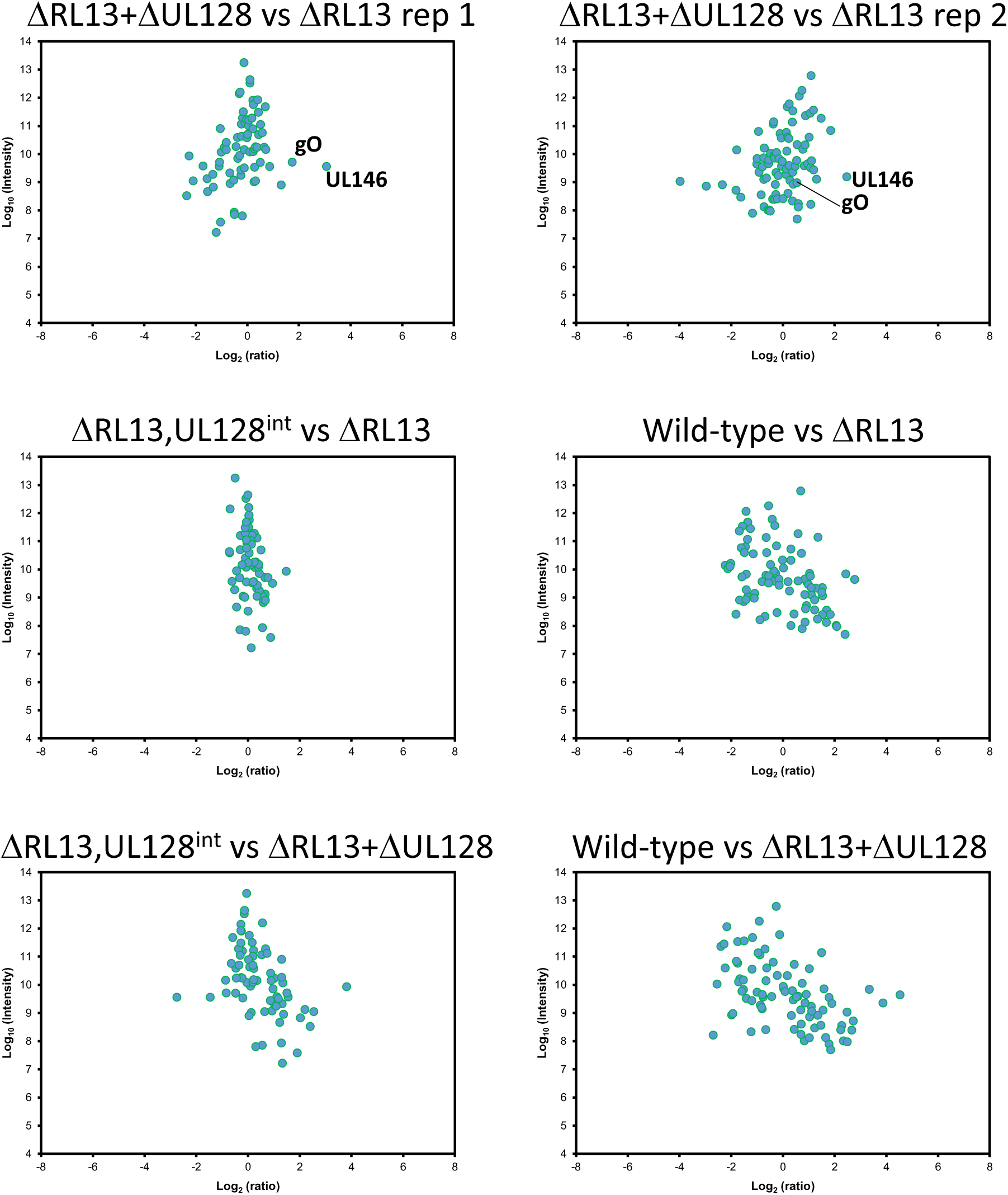
Scatter plot of viral proteins quantified in each SILAC-labelled virion experiment. The summed ion intensity (y axis) is shown as log10, and ratio (x-axis) as log2.

**Figure S2:**
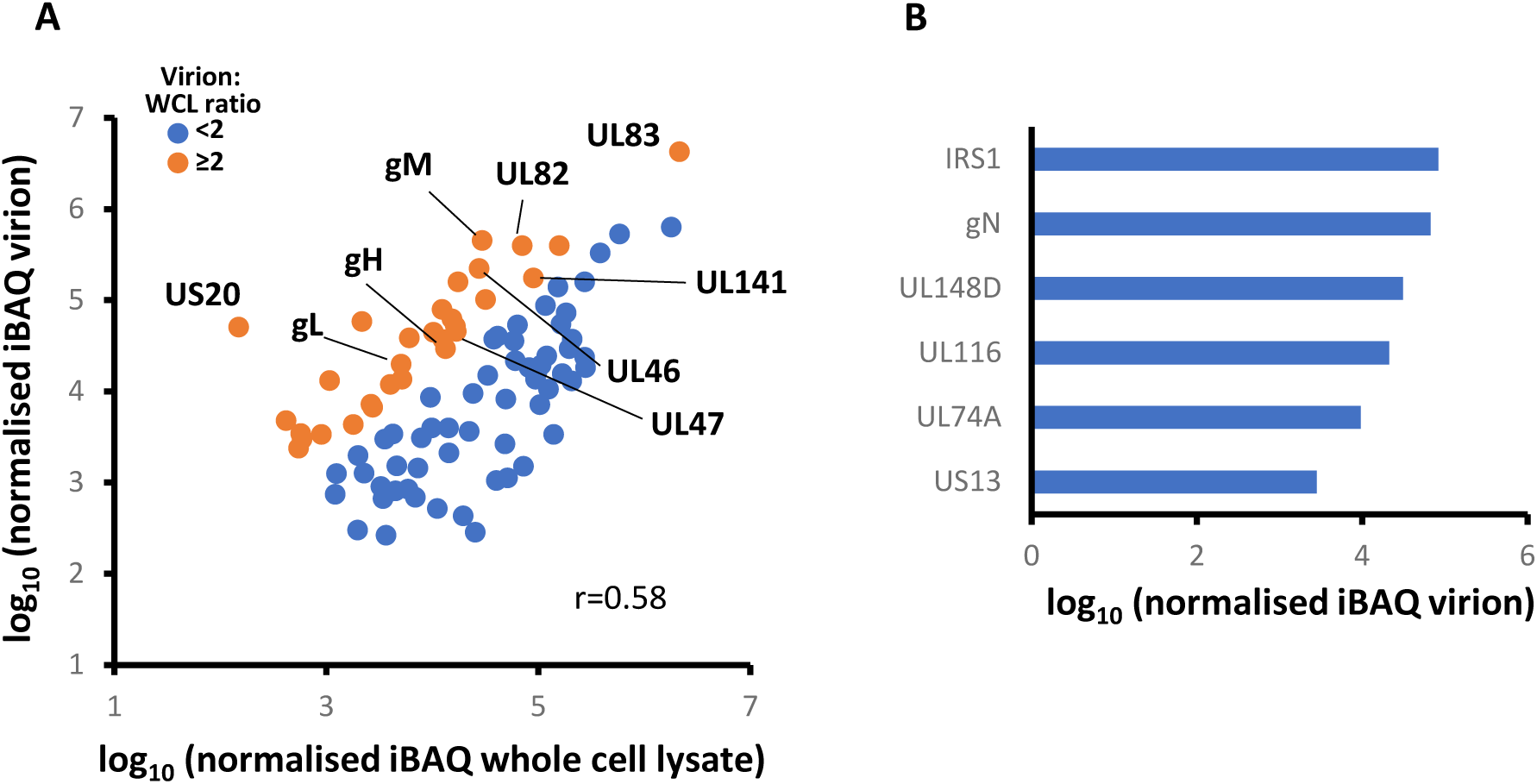
(A) comparison of abundance of HCMV proteins in the virion and in whole cell lysates (WCL) of infected cells. Proteins displayed in orange had a virion:WCL ratio of >2. (B) HCMV proteins only detected in the virion but not in WCL of infected cells. All quantified proteins are shown in Table S3.

**Figure S3:**
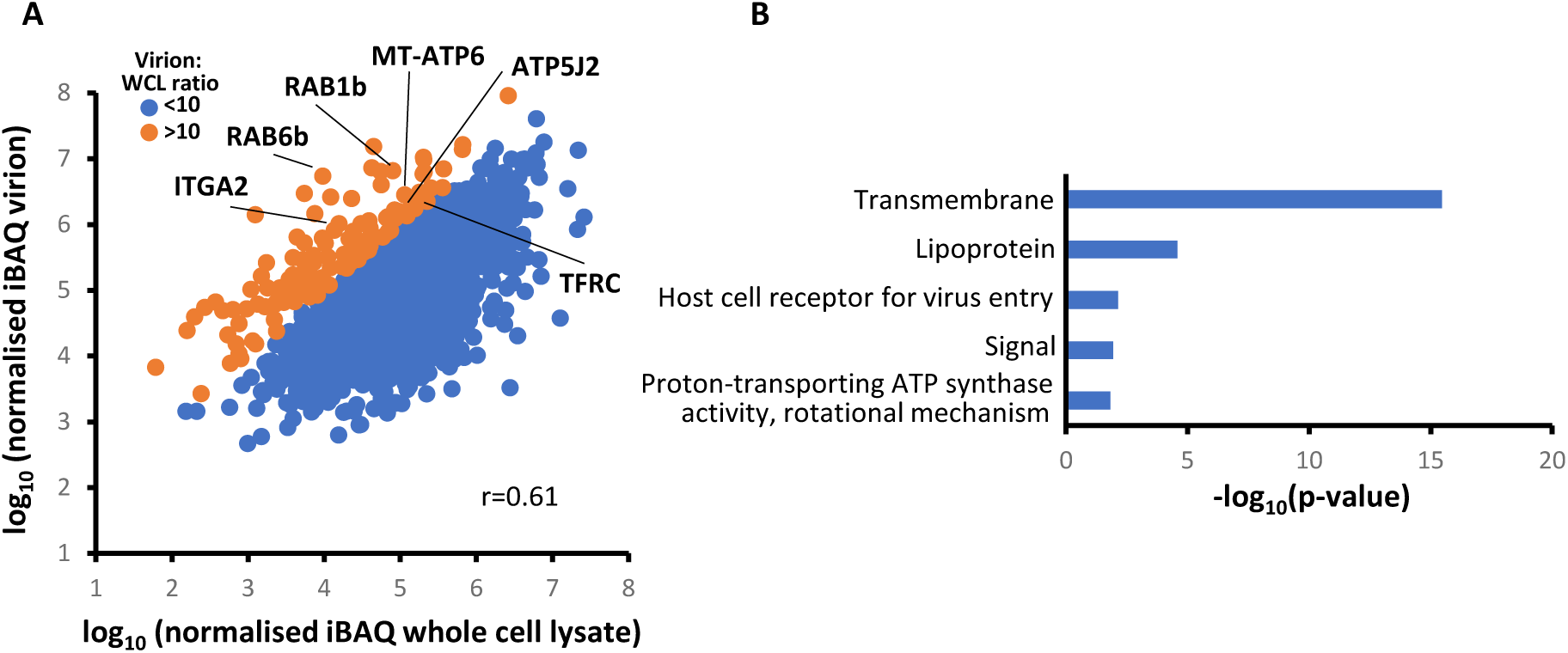
(A) comparison of abundance of human proteins in the virion and in whole cell lysates (WCL) of infected cells. Proteins displayed in orange had a virion:WCL ratio of >10. All quantified proteins are shown in Table S3B. (B) Enrichment of pathways within human proteins with a virion:WCL ratio of >2, using DAVID software, in comparison to all quantified human proteins. Full data are shown in Table S3C.

**Figure S4.**
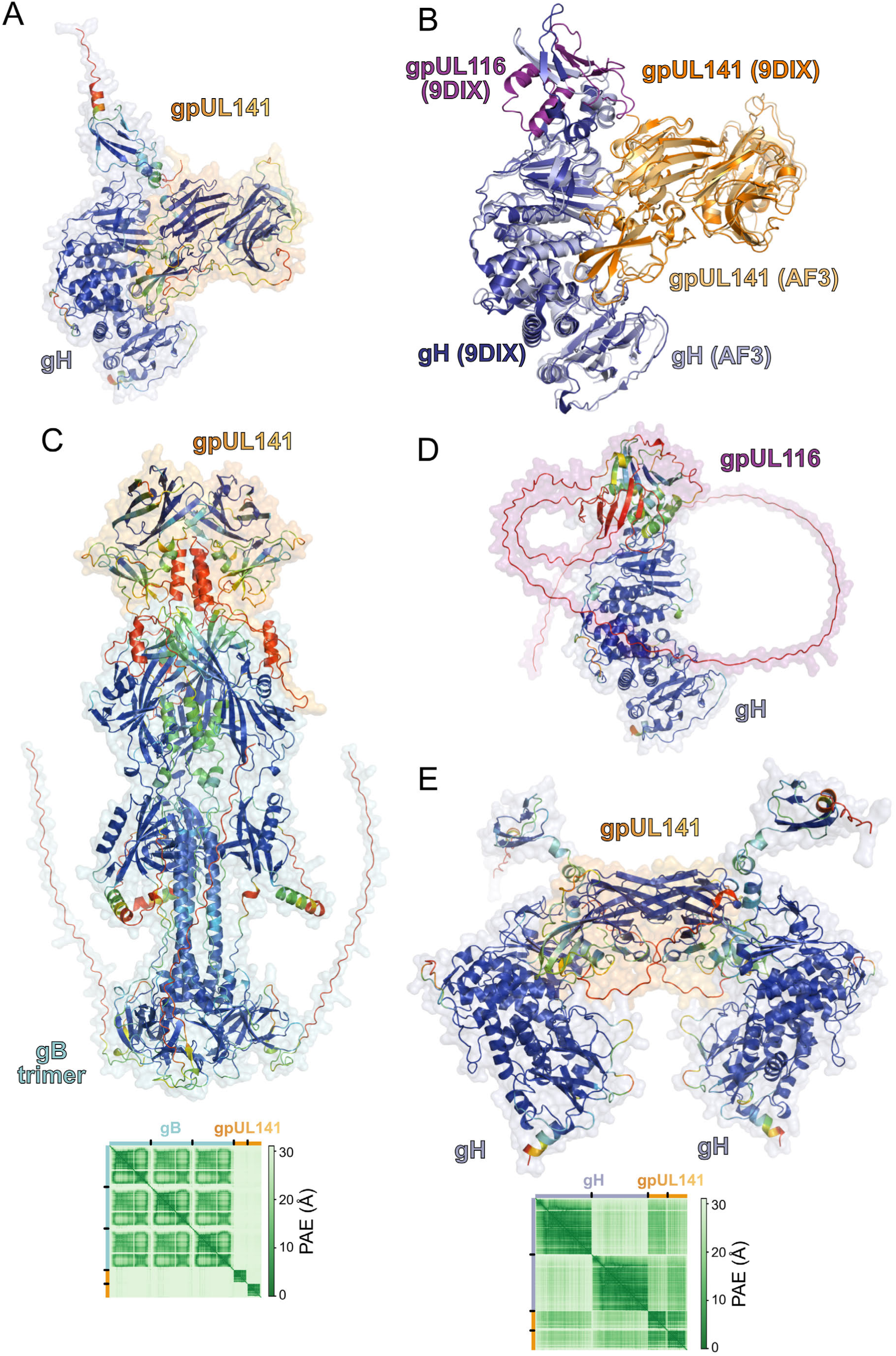
AlphaFold3 (AF3) predictions of HCMV glycoprotein complexes. (A) Predicted structure of gH (blue semi-transparent surface) in complex with a homodimer of gpUL141 (orange semi-transparent surface). Proteins are shown as ribbons coloured by prediction confidence (predicted local distance difference test; pLDDT), from red (low confidence, pLDDT ≤ 50) to blue (high confidence, pLDDT ≥ 90). (B) Superposition of the predicted gpUL141:gH complex (light orange and blue, respectively) onto the experimental structure (PDB 9DIX^46^) of gpUL141 in complex with gH and gpUL116 (dark orange, blue and purple, respectively). For clarity, residues that were not modelled in the experimental structure are omitted from the AF3 model. The AF3 model is highly similar to the experimental structure (root mean squared deviation of 1.8 Å across 960 Cα residues). (C) Predicted complex between a homotrimer of gB (cyan semi-transparent surface) and a homodimer of gpUL141 (orange semi-transparent surface), with ribbons coloured as in (A). gB has been predicted in the post-fusion conformation^83^. Inset shows the predicted aligned error (PAE) of the complex, indicating low confidence in the orientation of gpUL141 relative to gB. (D) Predicted complex between a gH (blue semi-transparent surface) a UL116 (purple semi-transparent surface), with ribbons coloured as in (A). (E) Predicted structure of two gH molecules (blue semi-transparent surfaces) in complex with a homodimer of gpUL141 (orange semi-transparent surface). The PAE plot (inset) indicates that the relative orientations of gH molecules with respect to the gpUL141 homodimer are predicted with high confidence.

