## Supplementary material for "Virion Proteomics of Genetically Intact HCMV Reveals a Novel Regulator of Envelope Glycoprotein Composition that Protects Against Humoral Immunity": Table S1

**Table S1. Virus encoded proteins identified by MS-analysis of Merlin virions.**

| ORF[a] | Rank[b] | ID[c] | Protein/Function[d] | AD169[e] | RhCMV[f] | MCMV[g] | TB40[h] | AD169-RV[i] |
|---|---|---|---|---|---|---|---|---|
| | | | **A- Proteins identified with high confidence** | | | | | |
| | | | ***Capsid:*** | | | | | |
| UL46 | 8 | 8/8 | pUL46/mCBP (minor capsid protein binding protein); interacts with mCP to form capsid triplex | X | X | X | X | X |
| UL48A | 17 | 8/8 | pUL48A/SCP (smallest capsid protein); decorates capsid hexons | X | X | X | X | X |
| UL80 | 39 | 8/8 | pUL80/PR-AP (assembly protein) precursor; scaffold for capsid assembly | X | X | X | X | X |
| UL85 | 6 | 8/8 | pUL85/mCP (minor capsid protein); interacts with mCBP to form capsid triplex | X | X | X | X | X |
| UL86 | 3 | 8/8 | pUL86/MCP (major capsid protein); forms capsid walls and faces | X | X | X | X | X |
| UL104 | 70 | 8/8 | pUL104/PORT; forms specialized portal capsomere for genome insertion/release | X | X | X | X | X |
| | | | ***Tegument:*** | | | | | |
| UL23 | 67 | 6/8 | Tegument. Inhibits transcription of IFN-γ stimulated genes | | | | | |
| UL24 | 19 | 8/8 | pUL24; US22 family member; enhances replication in endothelial cells | X | X | | X | X |
| UL25 | 7 | 8/8 | pUL25; UL25 family member | X | X | X | X | X |
| UL26 | 16 | 8/8 | pUL26; US22 family member, MIEP transcription transactivator, tegument material phosphorylation, stabilises virion | X | X | | | X |
| UL28/UL29 | 79 | 5/8 | pUL28/UL29; US22 family member, Stimulates IE gene expression | | | X | | X |
| UL32 | 12 | 8/8 | pp150 / NSP (nucleocapsid-proximal stabilization protein); virion maturation | X | X | X | X | X |
| UL35 | 41 | 8/8 | pUL35; UL25 family member, virion morphogenesis | X | X | X | X | X |
| UL36 | 50 | 7/8 | pUL36x1; US22 family member, vICA (viral inhibitor of caspase-8-induced apoptosis) | | | | | |
| UL43 | | 8/8 | pUL43; US22 family member | X | | X | | X |
| UL45 | 15 | 8/8 | pUL45/RR1 (Ribonucleotide reductase homologue sub-unit) | X | X | X | X | X |
| UL47 | 28 | 8/8 | pUL47/LTPbp (Largest tegument protein binding protein); interacts with ppUL48 intra-cellular transport of nucleocapsid | X | X | X | X | X |

| | | | | | | | | |
|---|---|---|---|---|---|---|---|---|
| UL48 | 33 | 8/8 | pUL48/LTP (Largest tegument protein); interacts with ppUL47 intra-cellular transport of nucleocapsid | X | X | X | X | X |
| UL50 | 38 | 8/8 | pUL50/NEC1 (nuclear egress complex membrane anchoring component 1); interacts with pUL53 (NEC2) to orchestrate nuclear egress of nucleocapsids | X | X | | | X |
| UL52 | 62 | 8/8 | pUL52; DNA encapsidation, nucleocapsid formation | | X | | | X |
| UL69 | 49 | 8/8 | ppUL69; MRP (multiple regulatory protein) | X | | X | X | X |
| UL71 | 47 | 8/8 | pUL71; Secondary envelopment, virion egress | X | X | X | X | X |
| UL82 | 5 | 8/8 | ppUL82/pp71/UMP (upper matrix protein); secondary envelopment, relieves DAXX-mediated repression | X | X | X | X | X |
| UL83 | 1 | 8/8 | ppUL83/pp65/LMP (lower matrix protein); phosphorylase, innate and adaptive immune response evasion | X | X | X | X | X |
| UL88 | 34 | 8/8 | pUL88; Putative cytoplasmic egress function | X | X | X | X | X |
| UL94 | 11 | 8/8 | pUL94; interacts with pp28 to facilitate secondary envelopment | X | X | X | X | X |
| UL96 | 76 | 7/8 | pUL96; interacts with pp150 to stabilise nucleocapsid during cytoplasmic egress | X | X | | | X |
| UL97 | 35 | 8/8 | ppUL97/VPK (viral protein kinase); phosphorylates viral and cellular proteins | X | X | X | X | X |
| UL99 | 20 | 8/8 | ppUL99/pp28; myristylated protein, secondary envelopment | X | X | X | X | X |
| UL103 | 54 | 7/8 | pUL103; VEP (Virion and DB egress protein); orchestrates release from producer cells | X | X | | | X |
| IRS1 | 18 | 8/8 | pIRS1; US22 family member, PKR (Protein kinase R) inhibitor, Transcriptional activator | X | | | | X |
| US22 | 32 | 8/8 | pUS22; US22 family member | X | X | X | | X |
| US23 | 91 | 8/8 | pUS23; US22 family member | X | | | | X |
| US24 | 82 | 7/8 | pUS24; US22 family member; enhances early infection | X | X | | | X |
| TRS1 | 85 | 8/8 | pTRS1; US22 family member, PKR (Protein kinase R) inhibitor, Transcriptional activator, capsid assembly | X | X | | | X |
| ***Transcription/replication machinery, yet to be assigned to virion compartment:*** | | | | | | | | |
| UL34 | 57 | 8/8 | pUL34; Represses US3 transcription | | | | | X |
| UL44 | 2 | 8/8 | ppUL44/PPS (Viral DNA polymerase processivity sub-unit); DNA synthesis | X | X | X | | X |
| UL57 | 55 | 8/8 | ppUL57/SSB (single-stranded DNA binding protein); DNA synthesis | X | | X | | X |
| UL70 | 73 | 8/8 | pUL70/HP2 (DNA helicase-primase sub-unit 2); DNA synthesis | | | X | | X |
| UL77 | 72 | 8/8 | pUL77/CVC1 (putative capsid vertex-specific component 1); DNA encapsidation | X | X | X | | X |

| | | | | | | | | |
|---|---|---|---|---|---|---|---|---|
| UL84 | 24 | 8/8 | ppUL84/Viral DNA replication accessory; DURP family member, DNA synthesis | X | X | | | X |
| UL89 | 60 | 8/8 | pUL89/TER1 (terminase sub-unit 1); DNA encapsidation | X | | | X | X |
| UL93 | 83 | 7/8 | pUL93/CVC2 (putative capsid vertex-specific component 2); DNA encapsidation | X | X | | X | X |
| UL98 | 44 | 8/8 | pUL98/NUC (deoxyribonuclease) | | X | X | | X |
| UL112-UL113 | 40 | 8/8 | pUL112/UL113; Orchestrates DNA synthesis, transcriptional activator | X | X | | | X |
| UL114 | 59 | 7/8 | ppUL114; DNA synthesis UNG (Uracil-DNA glycosylase), control DNA synthesis | | | | | X |
| UL122 | 48 | 8/8 | pUL122; IE2; transactivator or host-cell transcription machinery | X | X | | X | X |
| UL146 | 66 | 7/8 | gpUL146; vCXC chemokine homologue family, putative chemokine | | | | | |
| UL54 | 81 | 5/8 | pUL54; POL; catalytic DNA polymerase sub-unit | X | | X | | X |
| UL56 | 86 | 6/8 | pUL56; TER2 (terminase sub-unit 2); DNA encapsidation | X | | X | | X |
| UL102 | 87 | 5/8 | pUL102/HP3 (helicase primase sub-unit 3); DNA synthesis | | X | X | | X |
| UL147 | 69 | 6/8 | pUL147; vCXC-2 chemokine homologue family, putative chemokine | | | | | |
| ***Envelope:*** | | | | | | | | |
| RL10 | 29 | 8/8 | gpRL10 | X | | | X | X |
| RL11 | 51 | 8/8 | gpRL11/gp34; RL11 family member, IgG Fc binding | | | | | X |
| RL12 | 25 | 8/8 | gpRL12/gp95; RL11 family member, IgG Fc binding | | | | | X |
| RL13 | 23 | 2/2 | gpRL13; RL11 family member, putative IgG Fc binding, impedes replication in fibroblasts and epithelial cells | | X | | | |
| UL16 | 30 | 7/8 | gpUL16; membrane glycoprotein, NK cell evasion, blocks NKG2D ligands MICB, ULBP1 and ULBP2 | | | | | |
| UL33x1 | 43 | 8/8 | pUL33x1; GPCR homologue family member, constitutive signalling GPCR | X | X | | X | X |
| UL37 | 53 | 5/8 | Isoform vMIA of pUL37; multifunctional transmembrane protein that plays several key roles in viral replication | | | | | |
| UL41A | 31 | 7/8 | | X | X | | X | X |
| UL55 | 13 | 8/8 | gB; gCI subunit, virion binding and entry | X | X | X | X | X |
| UL73 | 21 | 8/8 | gN; gCII sub-unit, virion binding, progeny virion secondary envelopment | X | X | | | X |
| UL74 | 92 | 8/8 | gO; gCIII sub-unit, fibroblast cell entry, release of progeny virions from producer cells | X | X | X | | X |

| | | | | | | | | |
|---|---|---|---|---|---|---|---|---|
| UL74A | 58 | 6/8 | gpUL74A; putative membrane glycoprotein | | | | | X |
| UL75 | 37 | 8/8 | gH; gCIII/pentameric complex sub-unit, entry into cells | X | X | X | | X |
| UL78 | 74 | 8/8 | gpUL78; GPCR homologue family member; putative chemokine receptor | | | | | |
| UL100 | 87 | 8/8 | gM; gCII sub-unit, virion binding, progeny virion secondary envelopment | X | X | X | X | X |
| UL115 | 52 | 7/8 | gL; gCIII/pentameric complex sub-unit, entry into cells | X | X | | | X |
| UL116 | 42 | 8/8 | gpUL116; putative membrane glycoprotein | | X | | | X |
| UL118-UL119 | 78 | 8/8 | gpUL118-UL119/gp68; IgG Fc binding | X | X | | X | X |
| UL128 | 45 | 5/6 | gpUL128; pentameric complex, entry into non-fibroblasts | | | | X | X |
| UL130 | 63 | 5/6 | gpUL130; pentameric complex, entry into non-fibroblasts | | | | | |
| UL131A | 75 | 6/6 | gpUL131A; pentameric complex, entry into non-fibroblasts | | | | X | |
| UL132 | 10 | 8/8 | gpUL132; enhances replication in fibroblasts | X | X | | X | X |
| UL139 | 94 | 5/8 | gpUL139; putative membrane glycoprotein | | | | | |
| UL140 | 95 | 6/8 | pUL140; putative membrane protein | | | | | |
| UL141 | 9 | 7/7 | gpUL141; UL14 family member, NK evasion, down regulates CD155 and CD112 | | X | | | |
| UL148 | 14 | 7/8 | gpUL148, putative membrane glycoprotein | | | | | |
| US9 | 46 | 7/8 | gpUS9; US6 family member, membrane glycoprotein | | | | | |
| US12 | 65 | 8/8 | pUS12; US12 family member, 7TM protein | | X | | | X |
| US13 | 77 | 6/8 | pUS13; US12 family member, 7TM protein | | | | | |
| US14 | 80 | 8/8 | pUS14; US12 family member, 7TM protein | | | | | X |
| US20 | 27 | 8/8 | pUS20; US12 family member, 7TM protein | | | | | X |
| US27 | 26 | 8/8 | gpUS27; GPCR homologue family member, 7TM protein, enhances virion release | X | | | X | |
| US28 | 56 | 8/8 | gpUS28; GPCR homologue family member, 7TM protein, CC and CXC3 chemokine receptor | | X | | | X |
| ***Uncharacterised:*** | | | | | | | | |
| RL1 | 68 | 5/8 | pRL1; RL1 family member, Degrades the host antiviral factor SLFN11 via the cullin4-RING E3 ubiquitin ligase (CRL4) complex | | | | | |
| UL13 | 88 | 6/8 | pUL13; modulates mitochondrial ultrastructure | | X | | | |

| | | | | | | | | |
|---|---|---|---|---|---|---|---|---|
| UL31 | 90 | 7/8 | pUL31; DURP family member | | | X | | |
| UL95 | 89 | 7/8 | Participates in the expression of late viral mRNAs | | | X | | |
| UL135 | 97 | 5/8 | pUL135; reactivation from latency, actin remodelling | | | | | |
| UL145 | 22 | 7/8 | pUL145; recruit DDB1-containing ubiquitin ligases to induce proteasomal degradation of STAT2. | | | | | |
| UL148D | 36 | 6/8 | Inhibits ADAM17 function | | | | X | |
| UL150A | 71 | 6/8 | Putative secreted protein | | | | | |
| US1 | 84 | 7/8 | US1 family member | | X | | | X |
| US8 | 61 | 7/8 | gpUS8; Type I membrane glycoprotein | | | | | |
| US26 | 93 | 6/8 | pUS26; US22 family member | | X | | | |
| **B- Proteins identified with low confidence** | | | | | | | | |
| ***Transcription/replication machinery, immune evasions, yet assigned to virion compartment:*** | | | | | | | | |
| UL105 | | 4/8 | pUL105; HP1 (helicase-primase sub-unit 1), DNA synthesis | | | X | | |
| UL123 | | 4/8 | ppUL123/IE1 (major IE protein); promotes transcription cascade, enhances IE2 activation | | | | | X |
| ***Envelope:*** | | | | | | | | |
| UL15A | | 1/8 | Uncharacterised | | | | | |
| UL40 | | 2/8 | Loads peptides onto HLA-E | | | | | |
| US3 | | 4/8 | Retains MHC class I heterodimers in the endoplasmic reticulum | | | | | |
| US18 | | 4/8 | pUS18; US12 family member, 7TM protein | | X | | | |
| ***Uncharacterised*** | | | | | | | | |
| RL9A | | 1/8 | | | | | | |
| UL5 | | 2/8 | May play a role in rearrangement of cellular cytoskeleton towards an efficient viral assembly and spreading. | X | | | | |
| UL14 | | 4/8 | Uncharacterised | | | | | |
| UL17 | | 3/8 | | | | | | |
| UL22A | | 3/8 | Glycoprotein | X | | | | X |

| | | | | | | | |
|---|---|---|---|---|---|---|---|
| UL27 | 1/8 | Uncharacterised | | | | | |
| UL30 | 1/8 | Uncharacterised | | X | | | |
| UL49 | 4/8 | pUL49; subunit of the viral pre-initiation complex, regulates gene transcription | X | | | | |
| UL53 | 4/8 | pUL53 (NEC2) (nuclear egress complex membrane anchoring component 2); interacts with pUL50/NEC1 to orchestrate nuclear egress of nucleocapsids | | | | | X |
| UL72 | 2/8 | Deoxyuridine 5-triphosphate nucleotidohydrolase | X | | X | | |
| UL76 | 4/8 | pUL76; modulates gene expression | | | | | |
| UL117 | 1/8 | Plays a role in the inhibition of host DNA replication in the infected cell | | | | | X |
| UL136 | 1/8 | Plays a role in latency | | | | | |
| UL138 | 3/8 | Modulates the expression of several host cell surface receptors | | | | | |
| UL150 | 1/8 | Uncharacterised | | | | | |
| US30 | 1/8 | Uncharacterised | | | | | |
| US32 | 4/8 | Uncharacterised | | | | | |

[a] ORFs encoding proteins identified in Merlin virions.
[b] rank of protein based on abundance in the wildtype HCMV virion proteome, with 1 being the most abundant as assessed by IBAQ.
[c] identities depict the number of virion preparations in which proteins were detected.
[d] comments as to the biochemistry and function of proteins identified, as described in Field's Virology
[e] proteins previously identified during mass spectrometry analysis of HCMV Strain AD169 virions[28]
[f] homologous proteins identified during mass spectrometry analysis of RhCMV virions[36]
[g] homologous proteins identified during mass spectrometry analysis of MCMV virions[37]
[h] homologous proteins identified during mass spectrometry analysis of HCMV Strain TB40 virions[30]
[i] proteins previously identified during mass spectrometry analysis of HCMV Strain AD169 virions[29]
